# Leptin Resistance in the Ovary of Obese Mice Is Associated with Profound Changes in the Transcriptome of Cumulus Cells

**DOI:** 10.1101/729657

**Authors:** Karolina Wołodko, Edyta Walewska, Marek Adamowski, Juan Castillo-Fernandez, Gavin Kelsey, António Galvão

## Abstract

**Background/Aims:** Obesity is associated with infertility, decreased ovarian performance and lipotoxicity. However, little is known about the aetiology of these reproductive impairments. Here, we hypothesise that the majority of changes in ovarian physiology in diet-induced obesity (DIO) are a consequence of transcriptional changes downstream of altered leptin signalling. Therefore, we investigated the extent to which leptin signalling is altered in the ovary upon obesity with particular emphasis on effects on cumulus cells (CCs), the intimate functional companions of the oocyte. Furthermore, we used the pharmacological hyperleptinemic (LEPT) mouse model to compare transcriptional profiles to DIO.

**Methods:** Mice were subjected to DIO for 4 and 16 weeks (wk) and leptin treatment for 16 days, to study effects in the ovary in components of leptin signalling at the transcript and protein levels, using Western blot, Real-time PCR and immunostaining. Furthermore, we used low-cell RNA sequencing to characterise changes in the transcriptome of CCs in these models.

**Results:** In the DIO model, obesity led to establishment of ovarian leptin resistance after 16 wk high fat diet (HFD), as evidenced by increases in the feedback regulator suppressor of cytokine signalling 3 (SOCS3) and decreases in the positive effectors phosphorylation of tyrosine 985 of leptin receptor (ObRb-pTyr985) and Janus kinase 2 (pJAK2). Transcriptome analysis of the CCs revealed a complex response to DIO, with large numbers and distinct sets of genes deregulated at early and late stages of obesity; in addition, there was a striking correlation between body weight and global transcriptome profile of CCs. Further analysis indicated that the transcriptome profile in 4 wk HFD CCs resembled that of LEPT CCs, in the upregulation of cellular trafficking and impairment in cytoskeleton organisation. Conversely, after 16 wk HFD CCs showed expression changes indicative of augmented inflammatory responses, cell morphogenesis, and decreased metabolism and transport, mainly as a consequence of the physiological changes of obesity.

**Conclusion:** Obesity leads to ovarian leptin resistance and major time-dependent changes in gene expression in CCs, which in early obesity may be caused by increased leptin signalling in the ovary, whereas in late obesity are likely to be a consequence of metabolic changes taking place in the obese mother.

## INTRODUCTION

Obesity is considered one of the major public health challenges of modern times and has been linked to various comorbidities, such as metabolic syndrome, type 2 diabetes, cancer, stroke (1) and infertility (2). Obese women have increased risk of menstrual dysfunctions and anovulation, pregnancy complications, and poor reproductive outcome (3). In mouse models, obesity is characterised by lipid accumulation in the ovary and ensuing lipotoxicity (4) and oxidative stress (5). Nonetheless, the exact mechanisms underlying ovarian pathogenesis in the course of obesity remain uncharacterised.

Leptin is a cytokine secreted by the adipose tissue (adipokine) (6). Indeed, soon after the introduction to an obesogenic environment, large amounts of leptin can be found in the circulation (7), making this adipokine one of the early-onset obesity markers. Leptin controls food intake through its action at the central nervous system (8); however, as a pleiotropic adipokine, leptin contributes to the regulation of numerous processes in the body, such as immune response (9) or angiogenesis (10). Concerning the reproductive tract, leptin has been shown to control the neuroendocrine reproductive axis (11) and folliculogenesis (12). Furthermore, leptin has been linked to ovulation (13) and embryo development (14). Leptin is detected in most cell types in the murine ovary, with the highest staining intensity seen in the oocyte (15). Nevertheless, no previous consideration has been made whether leptin signalling is dysregulated in the obese ovary. The leptin receptor b (ObRb) is a type I cytokine receptor, which signals through the association with the tyrosine kinase Janus kinase 2 (JAK2). Upon leptin binding and dimerization of the receptor, JAK2 is recruited (16), mediating the phosphorylation of three conserved tyrosine residues on the intracellular domain of the receptor: tyrosine (Tyr) 985, Tyr1077 and Tyr1138. Subsequently, signalling molecules are recruited to these activated tyrosines (17) and, as a result, the signal transducer and activator of transcription (STAT) 5 and/or STAT3 are also phosphorylated and translocated into the nucleus, where they regulate transcription (8,18). During sustained activation of ObRb, the expression of both suppressor of cytokine signalling 3 (SOCS3) and tyrosine-protein phosphatase 1B (PTP1B) is initiated as a negative feedback response (19,20). While PTP1B dephosphorylates JAK2, SOCS3 binds to receptor domains within JAK2 and Tyr985 and terminates signal transduction.

To date, little is known about the integrity of leptin signalling in the ovary during obesity progression, and its particular impact on the cumulus oophorous complex (COC). Cumulus cells (CCs) are vital regulators of oocyte growth and metabolism (21), controlling meiosis resumption (22), as well as the ovulation process itself (23). Thus, a better knowledge of the pathophysiology of the events taking place in these cells in the course of obesity will allow us to understand the molecular mechanisms leading to impaired ovarian performance and infertility. Furthermore, the transcriptional signature of CCs has been used to predict oocyte competence or embryo quality (24,25), demonstrating the importance of such references in assisted reproduction techniques. Here, we first characterise the establishment of leptin resistance in whole ovaries from diet induced obesity (DIO) mice fed for 4 and 16 weeks (wk). Subsequently, we analysed the transcriptome of CCs throughout obesity progression and identify temporally altered gene expression signatures. Finally, using a mouse model for pharmacological hyperleptinemia (LEPT), we pinpoint the transcriptional events mediated by increased leptin signalling in CCs in the early stages of obesity.

## MATERIALS AND METHODS

### Animal protocol

Breeding pairs from mouse strains C57BL/6J (B6) and B6.Cg-Lep^ob^/J (*ob/ob*) were obtained from the Jacksons Laboratory (The Jacksons Laboratory, Bar Harbour, Maine, USA). At 8 wk of age B6 animals were divided into 2 groups (n=12/ group). In DIO, the control group was fed *ad libitum* chow diet (CD, 11% energy in kcal from fat, 5053, rodent diet 20, LabDiet IPS, London, UK), while the experimental group received high fat diet (HFD, 58% energy in kcal from fat, AIN-76A 9G03, LabDiet IPS). Mice were maintained on the respective diet for 4 or 16 wk. Regarding the pharmacological hyperleptinemia protocol, 8 wk old B6 female mice fed CD were divided into two groups (n=15/group): i) saline 16 days (d) (CONT); ii) leptin 16 d (Recombinant Mouse Leptin, GFM26, Cell Guidance Systems, Cambridge, UK). The animals were injected intraperitoneally twice a day, at 09:00 h and 21:00 h and total dosage of 100 μg/day of leptin was administrated. Concerning the *ob/ob* model, mice were kept from weaning until twelve wk of age on CD. For all protocols, mice were housed with a 12 h light/12 h dark cycle at room temperature (23□, RT).

For phenotype characterisation of DIO, changes in body composition were monitored every 4 wk by nuclear magnetic resonance (NMR, Bruker, Rheinstetten, Germany), whereas in the LEPT model animals were phenotyped every three days. Body weight (BW), fat mass (FM), lean mass (LM), adiposity index (AI, fat mass/lean mass), and food intake (FI) were measured. For DIO and LEPT models vaginal cytology was done at 9:00 h, for twelve consecutive days. After vaginal lavage with saline, the smears were placed on clean glass slides and stained with Diff Quick Staining Set (Diff- Quick Color Kit, Rapid Staining Set, Medion Grifols Diagnostics AG, Duedingen, Switzerland), for cell identification as previously described (26). The results were analysed for percent of time spent in oestrous stage (further explained in Figure supplement 1J, 1K; Figure supplement 6G). All samples for mRNA, protein analysis and staining were collected at the oestrus stage.

A superovulation protocol was used for COC collection and further isolation of a pure population of CCs after removing the metaphase II (MII) oocytes. This developmental state of granulosa cells (GC) was preferred to ensure a tightly controlled level of progression. The animals were injected with pregnant mare’s serum gonadotropin (PMSG, G4877, 5IU, Sigma Aldrich, Saint Louis, Missouri, USA) followed after 48 h by human chorionic gonadotropin (hCG, Chorulon, 5IU, MSD Animal Health, Boxmeer, Holland). Subsequently, 18 h after hCG injection the animals were sacrificed.

### Ovary collections

Mice were sacrificed by cervical dislocation and the reproductive tracts collected and rinsed with phosphate buffered saline (PBS, 0.1M, pH=7.4). Ovaries were then removed from the genitalia and cleaned of adipose tissue. Ovaries were stored either in TRI Reagent (T9424, Sigma Aldrich) for mRNA, or radioimmunoprecipitation assay buffer (RIPA, 89901, ThermoFisher Scientific, Waltham, Massachusetts, USA) supplemented with protease inhibitor cocktail (PIC, P8340, Sigma Aldrich), phenylmethylsulfonyl fluoride (PMSF, P7626, Sigma Aldrich) and phosphatase inhibitor (88667, Thermo Fisher Scientific), for protein analysis. Samples were stored in −80°C, except for the analysis of phosphorylated proteins, in which samples were isolated immediately after sacrificing the animals.

### Ovarian cell isolation protocol

For theca and stroma enriched (TC) fraction collection, immediately after culling the animals, ovaries were transferred to Dulbecco’s modified Eagle’s medium (DMEM, D/F medium; 1:1 (v/v), D-8900, Sigma Aldrich) with 3% bovine serum albumin (BSA, 735078, Roche Diagnostics GmbH, Mannheim, Germany), 20 µg/ml gentamicin (G1397, Sigma Aldrich) and 250 μg/ml amphotericin (A2942, Sigma Aldrich). For the TC fraction, ovaries were punctured with a 16 gauge needle as described before (27). Briefly, after removing GC and oocytes, the remaining tissue was washed twice with media and stored in TRI Reagent for mRNA analysis (n=8/group). For CCs, after superovulating the animals, COCs were retrieved from oviducts, and further digested with hyaluronidase (H3506, 400 μg/ml, Sigma Aldrich). After removing the oocytes, a total of approximately 50 pure CCs were collected from one individual animal from either 4 wk or 16 wk DIO (n=5/ condition), as well as the 16 d LEPT (n=at least 3/ condition) protocol.

### RNA isolation and cDNA synthesis

For mRNA extraction, either whole ovaries or TC fraction were collected from mice in oestrus stage, placed in 1 ml of TRI Reagent in 1.5 ml eppendorf tubes (n=8/ group) and mechanically disrupted with a lancet. The suspension was pipetted up and down vigorously and incubated for 5 minutes (min) at RT. After centrifugation (9400 g, 4□, 15 min), the supernatant was transferred to a fresh tube and thoroughly mixed with 100 μl of 1-Bromo-3-chloropropan (BCP, BP151, Molecular Research Centre, Cincinnati, Ohio, USA), followed by incubation at RT for 10 min. Subsequently, samples were centrifuged (13500 g, 4□, 15 min) and the aqueous phase transferred to a new tube, before being mixed with an equal volume of isopropanol and incubated at −80°C for 60 min. Another centrifugation (20000 g, 4□, 15 min) to pellet down the RNA, which was then washed three times with 75% ethanol and incubated overnight in −80°C. Next day, samples were centrifuged (20000 g, 4□, 15 min) and the RNA pellet dried and resuspended in 20 μl of RNAse free water (W4502, Sigma Aldrich), supplemented with RNAse Inhibitor (RiboProtect, RT35, BLIRT, Gdańsk Poland). Finally, RNA quality and concentration were assessed with NanoDrop. Absorbance ratio at 260 nm and 280 nm (A260/A280) was determined and the quality and concentration of isolated mRNA confirmed.

A total of 1 μg of RNA was reversely transcribed using Maxima First Strand cDNA Synthesis Kit for Real-time polymerase chain reaction (PCR) (K1642, ThermoScientific) according to the manufacturer’s instructions. The cDNA was stored in −20°C until the real-time PCR was carried out.

### Real-time polymerase chain reaction

Real-time PCR was performed in a 7900 Real-Time PCR System (Applied Biosystems, Warrington, UK) using Maxima SYBR Green/ROX qPCR Master Mix (K0223, ThermoScientific). Primers were designed using Primer 3.0 v.0.4.0. software (28), based on gene sequences from GeneBank (NCBI), as described before (29). All primers were synthesised by Sigma Aldrich. Primer sequences, expected PCR products length,and GeneBank accession numbers are reported in Table 1. The total reaction volume was 12 μl, containing 4 μl cDNA (10μg), 1 μl each forward and reverse primers (80 nM or 160 nM), and 6 μl SYBR Green PCR master mix. Real-time PCR was carried out as follows: initial denaturation (10 min at 95□), followed by 45 cycles of denaturation (15 s at 95□) and annealing (1 min at 60□). After each PCR, melting curves were obtained by stepwise increases in temperature from 60 to 95□ to ensure single product amplification. In each real-time assay, both the target gene and a housekeeping gene (HKG) - *Ribosomal Protein L37* (*Rpl37*, primers in Table 1) or *Eukaryotic Translation Initiation Factor 5A* (*Eif5a*, primers in Table 1) - were run simultaneously and reactions were carried out in duplicate wells in a 384-well optical reaction plate (4306737, Applied Biosystems). The HKG selection was performed with NormFinder, in each experimental group. Real-time PCR results were analysed with the Real-time PCR Miner algorithm (30).

### Western blot

Protein expression was assessed by western blot (n=5/group). Ovaries from mice in oestrus stage were collected into RIPA supplemented with inhibitors and mechanically disrupted with a lancet. Then, lysates were incubated for one hour on ice, with mixing every 15 min. Subsequently, samples were centrifuged (20000 g, 4□ 15min) and the supernatant was collected and stored in −80°C until the analysis. The protein concentration was assessed using bicinchoninic acid assay (BCA, BCA1-1KT, Sigma Aldrich). A total of 10-40 μg of protein was loaded on 8-14% acrylamide gel, and after electrophoresis proteins were transferred to polyvinylidene difluoride (PVDF) or nitrocellulose membrane depending on the antibodies to be used. Membranes were blocked in 5% BSA (A2153, Sigma Aldrich) and incubated with primary antibodies (AB) overnight at 4°C. Leptin receptor and its phosphorylated domains were evaluated using the following antibodies: mouse monoclonal (MM) against leptin receptor (ObR; 1:500, cat# sc-8391, Santa Cruz Biotechnology, Dallas, Texas, USA), goat polyclonal (GP) against phosphorylated Tyr 985 ObRb (pTyr985ObRb; 1:500, cat# sc-16419, Santa Cruz Biotechnology), rabbit polyclonal (RP) against phosphorylated Tyr 1077 ObRb (pTyr1077ObR; 1:500, cat# 07-1317, Merck Millipore, Burlington, Vermont, USA), GP against phosphorylated Tyr 1138 ObRb (pTyr1138ObRb; 1:500, cat# sc-16421, Santa Cruz Biotechnology). The expression of other leptin signalling pathway components was assessed using the following antibodies: RP against JAK2 (1:200, cat# sc-294, Santa Cruz Biotechnology), RP against phosphorylated Tyr 1007/1008 JAK2 (pJAK2; 1:200, cat# sc-16566-R, Santa Cruz Biotechnology), RP against STAT3 (1:200, cat# sc-482, Santa Cruz Biotechnology), MM against phosphorylated Tyr 705 pSTAT3 (1:200, cat# sc-8059, Santa Cruz Biotechnology), RP against STAT5 (1:200, cat# sc-835, Santa Cruz Biotechnology), MM against phosphorylated Tyr 694/699 pSTAT5 (1:200, cat# sc-81524, Santa Cruz Biotechnology), GP against PTP1B (1:200, cat# sc-1718, Santa Cruz Biotechnology), MM against SOCS3 (1:500, cat# sc-51699, Santa Cruz Biotechnology) in cell lysates. The results were normalized with β-actin (1:10000, MM, cat# A2228, Sigma-Aldrich). All antibodies specifications are summarised in Table 2. Proteins were detected after incubation of the membranes with secondary GP anti-rabbit alkaline phosphatase-conjugated antibody (1:30000, cat# A3687, Sigma Aldrich), GP anti-mouse alkaline phosphatase-conjugated antibody (1:20000, cat# 31321, ThermoScientific), RP anti-goat alkaline phosphatase-conjugated (1:30000, cat# A4187, Sigma Aldrich), or RP anti-goat horseradish peroxidase-conjugated antibody (1:75000, cat# A50-100P, Bethyl, Montgomery, Alabama, USA) for 2 h at RT. Immune complexes were visualized using the alkaline phosphatase visualization procedure or ECL substrate visualization. Blots were scanned in a Molecular Imager VersaDoc MP 4000 System (BioRad, Hercules, California, USA) and specific bands quantified using ImageLab Software (BioRad). Finally, band density for each protein was normalised against β-actin.

### Immunohistochemistry and immunofluorescent staining

Ovaries collected from mice in oestrus stage (n=3/group) were fixed in 4% neutral phosphate-buffered formalin (NBF, 432173427, Poch, Gliwice, Poland) at 4°C for 24 h, and subsequently dehydrated in ethanol. Paraffin embedded ovarian tissues were sectioned into 5 μm slices. For antigen retrieval, sections were heated in citrate buffer (10 mM, pH=6.0). Tissue was incubated in blocking solution (ab64261, Abcam, Cambridge, UK) for 1 h at RT and primary RP anti-SOCS3 antibody (1:1000, ab16030, Abcam) or primary RP anti-PTP1B antibody (1:500, ab189179, Abcam) added overnight at 4°C. The negative control sections were incubated with RP anti-immunoglobulin G (IgG, cat# ab37415, Abcam) or without primary antibody. The primary antibody complexes were detected after incubating the tissue with biotinylated goat anti-rabbit IgG (H+L) (ab64261, Abcam) for 60 min, and streptavidin peroxidase for 40 min. Staining was evident after 15 s incubation in 3,3-diaminobenzidine (DAB) peroxidase substrate solution (Rabbit-specific HRP/DAB (ABC) Detection IHC Kit, ab64261, Abcam). Subsequently, samples were counterstained with haematoxylin (MHS16, Sigma Aldrich) and mounted. Sections were examined using Axio Observer Systems Z1 microscope (Carl Zeiss Microscopy GmbH, Hannover, Germany) and Zeiss ZEN 2.5 lite Microscope Software (Carl Zeiss, Germany). For immunofluorescence (IF), 5 μm sections were deparaffinised and rehydrated in an ethanol series. Next, tissues were permeabilised in 0.3% Triton X-100 (T8787, Sigma Aldrich), followed by antigen retrieval in citrate buffer (10 mM, pH=6.0) for 40 min in 90□ and blocking in 2% BSA (A2153, Sigma Aldrich) with 0,3 M glycine (G8898, Sigma Aldrich) in PBS-0.1% Tween 20 (P7949, Sigma Aldrich) (PBST) solution. Sections were then incubated with 0.3% Sudan Black (199664, Sigma Aldrich) in 70% ethanol for 10 min at RT, followed by washes in PBST. Slides were subsequently incubated with RP anti-SOCS3 antibody (1:200, ab16030, Abcam) overnight at 4°C. The negative control sections were incubated with RP anti-IgG (1:200) as before, or without primary antibody. On the next day slides were washed in PBST, followed by incubation with cyanine 3 (Cy3)-donkey polyclonal anti-rabbit IgG (H+L) (711-165-152, Jackson ImmunoReserach, Cambridgeshire, UK), and a series of washes in PBST. Finally, slides were covered with a drop of Prolong Gold medium with diamidino-2-phenylindole (DAPI) and sealed with cover slips. Images were captured using 40x/1.2A or 63x/1.4A oil immersion objectives on a LSM800 confocal microscope (Carl Zeiss, Germany).

### Enzyme-linked immunosorbent assay

Animals in oestrus were sacrificed (n=8/group) and blood samples collected after puncturing the heart. Blood samples were centrifuged (180 g, 4□ 10 min) and plasma stored at −80□. Levels of circulating leptin and insulin were assessed with enzyme-linked immunosorbent assay (ELISA) kit, according to the manufacturer’s instructions (Mouse Leptin ELISA Kit, 90030; Crystal Chem, Zaandam, Netherlands; Rat/Mouse Insulin ELISA Kit, cat. EZRMI-13K; Merck Millipore). The intra- and interassay coefficients of variation (CVs) were as follows: for Leptin ELISA kit <10% both and for Insulin ELISA kit 8.35% and 17.9%, respectively. To determine SOCS3 in ovarian extracts, ELISA test was used (ELISA KIT for SOCS3; cat no SEB684Mu, Cloud-Clone, Texas, USA). Briefly, the tissue was minced in lysis buffer (n=8/group), centrifuged (10000 g, 4□, 5 min) and protein concentration in the lysate determined with BCA test. All tests and assessments were performed according to the manufacturer’s instructions.

### RNA-seq library generation

Once the 16 wk HFD group presented divergence in BW gain, we identified 3 animals with less than 33 g of body weight that we designated as HFD low gainers (HFDLG) and excluded them from the regular HFD group for the further description of differently expressed genes (DEGs) between CD and HFD. CCs were collected into RLT buffer (1053393, Qiagen, Hilden, Germany) and kept at −80°C until library generation. Subsequently, RNA sequencing (RNA-seq) libraries were prepared following a previously described protocol (31,32), with minor changes. Briefly, mRNA was captured using Smart-seq2 oligo-dT pre-annealed to magnetic beads (MyOne C1, Invitrogen, Carlsbad, California, USA). The beads were resuspended in 10 μl of reverse transcriptase mix (100□U, SuperScript II, Invitrogen; 10 U, RNAsin, Promega, Madison, Wisconsin, USA), 1□×□Superscript II First-Strand Buffer, 2.5 mM ditiotreitol (DTT, Invitrogen), 1 M betaine (Sigma Aldrich), 9 mM magnesium chloride (MgCl_2_, Invitrogen), 1 μ Template-Switching Oligo (Exiqon, Vedbaek, Denmark), 1 mM deoxyribonucleotide triphosphate (dNTP) mix (Roche) and incubated for 60 min at 42°C followed by 30 min at 50°C and 10 min at 60°C (31,32). Amplification of the cDNA was then undertaken after adding 11 μl of 2□×□KAPA HiFi HotStart ReadyMix and 1 μl of 2 μ ISPCR primer (31,32), followed by the cycle: 98°C for 3 min, then 9 cycles of 98°C for 15 s, 67°C for 20 s, 72°C for 6 min and finally 72°C for 5 min. Finally, the cDNA was purified using a similar volume of AMPure beads (Beckman Coulter, Brea, California, USA) and eluted into 20 μl of nuclease- free water (P1195; Promega). All libraries were prepared from 100 to 200 pg of cDNA using the Nextera XT Kit (Illumina, San Diego, California, USA), according to the manufacturer’s instructions. The final cDNA libraries were purified using a 0.7:1 volumetric ratio of AMPure beads before pooling and sequencing on an Illumina Nextseq500 instrument in 75-base pair (bp) single-read high output mode at the Babraham Institute Sequencing Facility. A total of 5-10 million mappable reads per sample were obtained, with an average of 30-50 million reads per condition.

### Library mapping and trimming

Trim Galore v0.4.2 was used with default parameters on raw Fastq sequence files. Mapping of the RNA-seq data was done with Hisat v2.0.5 against the mouse GRCm38 genome, as guided by known splice sites taken from Ensemble v68.

### RNA-seq differential expression analysis

Mapped RNA-seq reads were quantified and analysed using SeqMonk version v1.45.4 (http://www.bioinformatics.babraham.ac.uk/projects/seqmonk/). Differential expression analysis was performed using DESeq2 (33) implemented in SeqMonk setting a false discovery rate (FDR) < 0.05.

### Statistical analysis

Statistical analysis was performed using GraphPad Prism 7.0. The D’Agostino-Pearson omnibus normality test was performed followed by nonparametric Mann-Whitney test or multiple unpaired t-test with statistical significance determined using the Bonferroni-Sidak method, depending on the experiment. The data are shown as the mean ± SD of three or more independent replicates. Significance was defined as values of p < 0.05.

### Statement of Ethics

All experiments were approved by Local Committee for the Ethical Treatment of Experimental Animals of Warmia- Mazury University (Agreement No. 80/2015, 15/2018, 38/2018), Olsztyn, Poland and were performed accordingly to the Guide for Care and Use of Laboratory Animals, endorsed by European legislation.

## RESULTS

### 1. Leptin signalling is impaired in the ovary of diet-induced obese mice

Initially we sought to characterise changes in leptin signalling in the ovary throughout DIO. Thus, mice were subjected to HFD for 4 and 16 wk and whole ovaries and TC fraction were collected for mRNA or protein analysis (Figure 1A). Oestrous stage was followed for twelve consecutive days, confirming that samples were collected in oestrus (Figure supplement 1J, 1K) in cycling animals. Mice significantly gained BW and FM already at 4 wk, with an average absolute gain in BW of 13 grams (g) in the 16 wk HFD group (Figure supplement 1A; p<0.0001). Three animals with comparable FI but BW gain less than 13 g were excluded from the statistical analysis and designated as HFD-low gainers (HFDLG). Also, after 4 and 16 wk HFD we confirmed high plasma levels of insulin (Figure supplement 1F; p<.0.01, p<0.001 respectively) and leptin (Figure supplement 1G; p< p<0.01, p<0.001 respectively) and the establishment of impaired glucose tolerance and insulin resistance at 16 wk HFD (Figure supplement 1H, 1I; p<0.01, p<0.001 respectively). Monitoring oestrous cycle revealed that the 4 wk HFD group had higher prevalence of oestrus counts compared with controls fed CD, whereas in the 16 wk HFD group there was a reduction in pro-oestrus (Figure supplement 1J, 1K; p<0.05, p<0.01 respectively).

**Figure 1.**
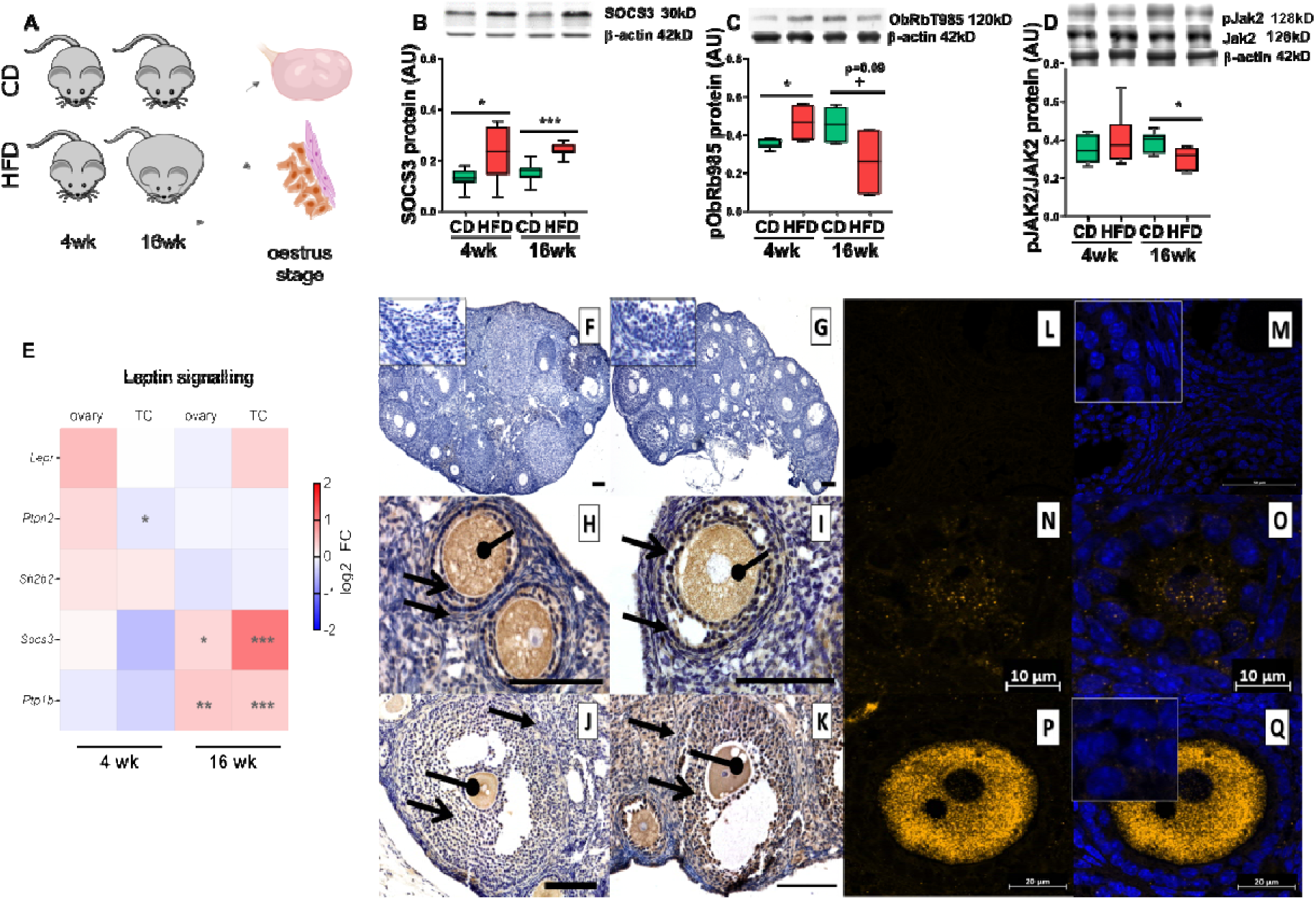
The establishment of leptin resistance in the ovary of diet induced obese mice. (A) Experimental design: animals were maintained on chow diet (CD) or high fat diet (HFD) for 4 weeks (wk) or 16 wk. Western blots with quantification, showing (B) total protein for SOCS3 (C) phosphorylation of tyrosine 985 of leptin receptor and (D) phosphorylation of Janus kinase 2. (E) Heatmap showing fold change in expression of mRNA of leptin signalling components measured in whole ovary or theca and stroma enriched (TC) fraction determined by RT-PCR. Immunohistochemical localisation of SOCS3 protein during follicle development in ovaries of mice subjected to diet-induced obesity (4 wk and 16 wk). Positive staining in brown, counterstaining with heamatoxylin. Negative control stained with polyclonal rabbit IgG (F) 4 wk CD and (G) 4wk HFD, localisation of SOCS3 in primary follicle (H) 4 wk CD and (I) 4 wk HFD, antral follicles (J) 16 wk CD and (K) 16 wk HFD. Staining is present in oocyte, granulosa and theca cells. Oval-headed arrow indicates oocyte; large-headed arrow indicates granulosa cells and small-headed arrow indicates theca cells. Scale bars represent 100µm. The staining was confirmed by immunofluorescent localisation of SOCS3 (L-Q). Positive staining in orange, nuclear counterstaining with DAPI in blue. (L-M) negative control 16 wk CD performed with polyclonal rabbit IgG, SOCS3 localised in (N-O) primordial follicles 16 wk CD, (P-Q) primary follicles 16 wk CD. Images are representatives of 3 biological replicates. Inserts in left top corners are magnifications of granulosa cells. mRNA expression of *Rpl37* and protein expression of β-actin were used to normalize the expression data. Each bar represents the mean ± SD. Differences between groups were analysed by Mann-Whitney test. N=4-8 for immunoblots and N=8 for RT-PCR analysis. * p<0.05; ** p<0.01; ***p<0.001; + p=0.09.

Next, we isolated protein from whole ovaries and studied the abundance of components of the leptin signalling pathway. Whilst we found initial hyperactivation of leptin signalling pathway as demonstrated by upregulation of SOCS3 protein (Figure 1B; Figure Supplement 2H; p<0.05 both) and a tendency to increased phosphorylation of STAT3 (Figure supplement 2E; p=0.06), after 16 wk HFD local leptin resistance was clearly established. This was evidenced by the decrease in abundance of leptin receptor (Figure supplement 2A; p<0.01), and a trend towards decreased phosphorylation of pTyr985 ObRb (Figure 1C; p=0.09) and decreased phosphorylation of JAK2 (Figure 1D; p<0.05), along with upregulation of SOCS3 (Figure 1B; p<0.001, Figure supplement 2H; p<0.05). In contrast, no differences were found in phosphorylation of other Tyr residues of ObRb (Figure supplement 2C and 2D) or in PTP1B expression (Figure supplement 2G). Additionally, there was reduced phosphorylation of STAT5 after 4 and 16 wk HFD (Figure supplement 2F; both p<0.01).

Next, we sought to characterise the extent to which various ovarian components responded similarly to increased circulating leptin during obesity. We performed real-time PCR analysis of whole ovaries and TC fraction. Despite no significant changes after 4 wk HFD, the mRNA of *Socs3* was increased after 16 wk HFD in both whole ovary (Figure 1E; p<0.05) and TC (Figure 1E; p<0.001), in comparison to the CD group. Additionally, the mRNA level of *Ptp1b* was increased in both whole ovary and TC after 16 wk HFD (Figure 1E; p<0.01, p<0.001 respectively).

The aforementioned results suggested that SOCS3 could be an important player in the establishment of leptin resistance in the ovary of obese mice. Therefore, we examined SOCS3 localisation in ovaries of DIO and the genetically obese model: mice with a mutation in the obese gene *(ob/ob)*. Immunohistochemistry (IHC) revealed the presence of SOCS3 protein in oocytes from follicles in all developmental stages (Figure 1H-K; Figure supplement 3E, 3F, 3I, 3J); in addition, theca cells and GC from the respective follicles were stained, as well as the ovarian stroma (Figure 1H-K). Importantly, we compared the IHC staining of SOCS3 and PTP1B in 16 wk HFD ovaries, which suggested that SOCS3 is the major ObRb inhibitor being expressed in the oocyte and GC, as PTP1B protein presented almost no staining in the oocyte (Figure supplement 3G, 3H, 3K, 3L). As a control for the specificity of the IHC data, we used confocal microscopy for immunofluorescence detection of SOCS3 on sections from DIO and leptin-deficient *ob/ob* (-/-) ovaries, confirming the localisation of SOCS3 (Figure 1N-Q) and observing a weaker intensity of SOCS3 in both oocyte and GC in ovaries from *ob/ob* (Figure supplement 3T, 3V) compared to wild type (+/+) (Figure supplement 3S, 3U). These IF results confirmed the specificity of the staining, since SOCS3 is expected to be less abundant in tissues form the *ob/ob* mouse (34). Furthermore, we also inferred that impaired leptin signalling in the ovary is likely to have direct implications for the oocyte, since the gamete was shown to express SOCS3.

### 2. Cumulus cell transcriptome analysis: global transcriptome of CCs reflects body weight

Next, we repeated the protocol and subjected the animals to superovulation in order to collect CCs and analyse the transcriptome from 4 wk and 16 wk DIO protocols (Figure 2A). A total of 50-80 CCs per animal were collected, from which RNA-seq libraries were generated using a Smart-seq2 oligo-dT method (31,32), with separate RNA-seq libraries made from the CC from each female. We then used Principal Component Analysis (PCA) to study the distribution of our samples according to global gene expression profile, and found that principal component 1 (PC1) was mainly driven by BW (Figure 2B). Here we decided to include the HFDLG from the 16wk HFD group as a control, to test whether the transcriptional response could be linked to the BW of the animals; indeed, the HFDLG samples clustered together with 16 wk CD of a similar weight (Figure 2B). The correlation between PC1 and BW was r=0.777 (p=3.026e-06) (Figure 2C; table 4), which substantiates the physiological effect driven by BW, rather than the nature of the diet itself, on the global gene expression profile of CCs.

**Figure 2.**
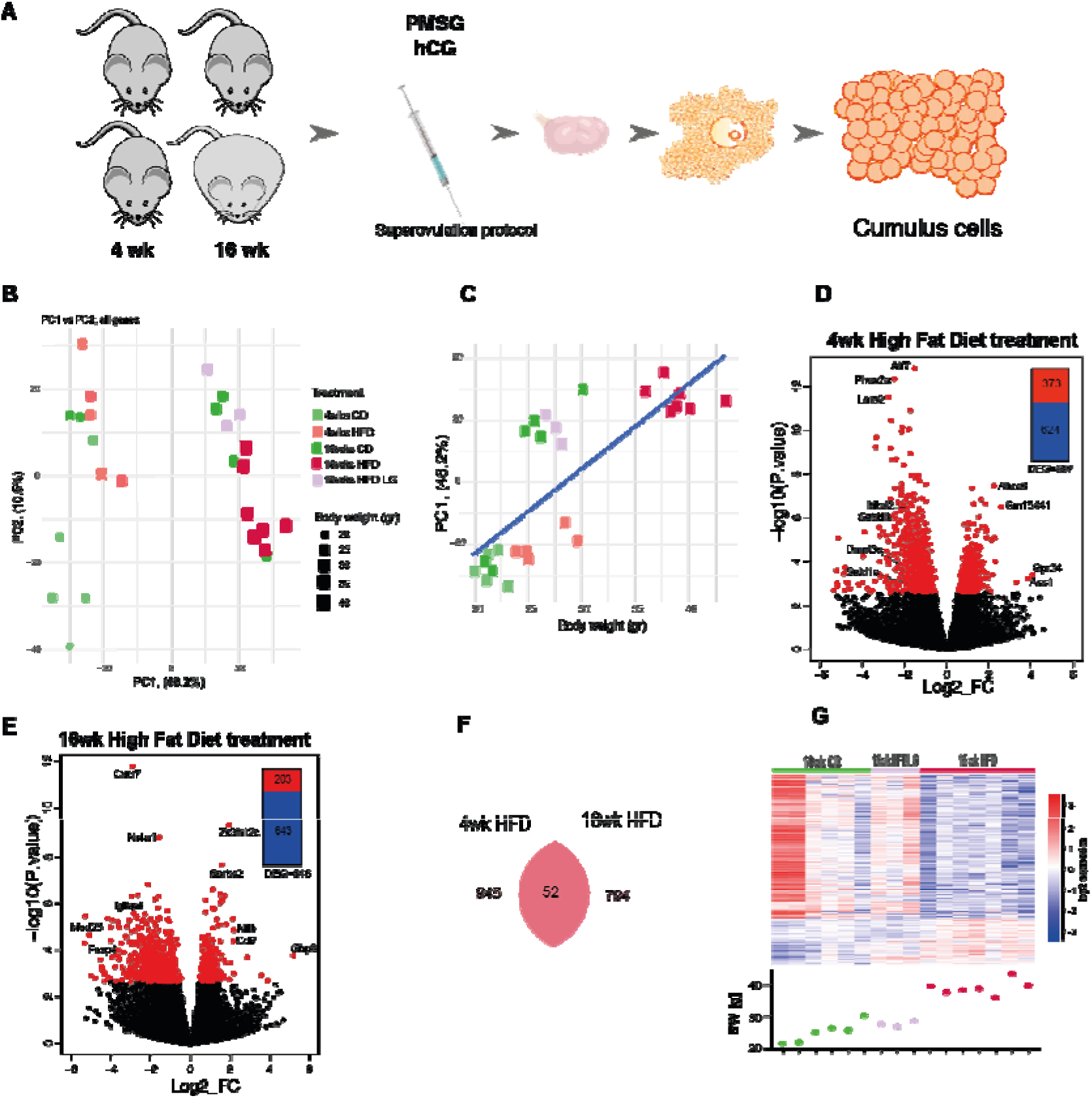
Cumulus cell transcriptome analysis in diet induced-obese mice reveals strong correlation with body weight. (A) Experimental design: mice were subjected to the indicated dietary protocol, superovulated and cumulus cells were collected from cumulus-oophorus-complexes. RNA-seq analysis of gene expression in cumulus cells obtained from mice after 4 or 16 weeks (wk) on chow diet (CD), high fat diet (HFD) or low gainers on HFD (HFDLG). N= 3-7 mice per group. (B) Principal component analysis of global transcriptome shows samples cluster into 2 groups accordingly to their body weight (BW). (C) Correlation of Principal Component 1 (PC1) with BW; r=0.777, p=3.026e-06. Volcano plots showing distribution of differentially expressed genes in (D) 4wks HFD and (E) 16wks HFD; genes with False Discovery Rate <0.05 coloured red. (F) Venn diagram showing the number of genes differentially expressed at false discovery rate (FDR) <0.05 between 4 wk and 16 wk groups. (G) Heatmap of 846 DEGs identified by DESeq2 analysis between 16wk CD and HFD CCs, including data for HFDLG CC samples, with BW of mice at time of collection plotted below. Heatmap representing fold of change of gene expression. log2_FC of reads per million (RPM).

Next, we aimed to identify DEGs in CCs: for this analysis, we excluded the 3 HFDLG outliers from the 16wk HFD, so as to ensure a minimum of 13 g of BW difference between CD 16 wk and HFD 16 wk and a BW difference of 5 g between CD 4 wk and HFD 4 wk (Figure Supplement 1A). After DESeq2 analysis (FDR <0.05), a total of 997 DEGs in 4 wk HFD (373 upregulated and 624 downregulated; Figure 2D; table 5) and 846 DEGs in 16 wk HFD (203 upregulated and 643 downregulated; Figure 2E table 5) were identified. Surprisingly, amongst the DEGs only 52 genes were common between the 4 wk and 16 wk comparisons (Figure 2F), highlighting the differences in pathophysiology of early and late stages of obesity. Gene ontology (GO) (35,36) analysis of the DEG lists showed that transcripts with increased abundance in 4 wk HFD were primarily linked to nitrogen and lipid metabolism and transport, but also cell stress and reactive oxygen species generation (Table 6). Transcripts downregulated after 4 wk HFD were mapped to pathways involved in regulation of macromolecule biosynthesis and gene expression, as well as chromatin organisation/histone modification and regulation of cell cycle (Figure supplement 4A; Table 6). After 16 wk HFD treatment, upregulated genes were associated with negative regulation of development and cellular component organisation, while pathways highlighted for downregulated genes included localisation, transport and positive regulation of metabolism (Figure supplement 4B; Table 7). Therefore, in this analysis we identified the gene signatures in CCs altered at the onset and later development of DIO.

Finally, we wished to examine the impact of BW as a factor on gene expression in CCs. Therefore, we examined the expression of the 846 DEGs identified in 16 wk HFD group in the 16 wk HFDLG and CD samples. Strikingly, for this set of genes, the HFDLGs presented an expression pattern closer to 16 wk CD than to 16 wk HFD (Figure 2G), revealing a very strong correlation between BW and global gene expression profile in CCs indicative of the impact of female physiology on CCs gene expression. As CCs represent an important accessible source of biomarkers for the assessment of reproductive potential of the mother, they are often sampled in assisted reproductive technologies (ART) to profile biomarkers of oocyte competence or embryo quality. Thus, we looked for known markers of embryo quality (25) in the 16 wk DEGs and discovered that *Nfib* was upregulated and *Ptgs2* and *Trim28* transcripts were downregulated in CCs (Figure supplement 5A-C). The altered expression of these markers in CCs during late obesity might indicate direct consequences for oocyte and embryo quality, as previously proposed (24,37–39).

### 3. Differential effects on gene expression in CCs early and late in obesity

We next sought to characterise how gene expression in CCs changes between the early and late stages of DIO. To do this, we first evaluated the expression of the 997 DEGs from 4 wk and the 846 DEGs from 16 wk HFD at both time-points, aiming to identify the directionality of gene signatures throughout obesity (Figure 3A). Note that in this analysis, only a minority of the DEGs are significantly altered at both time points (as noted above in figure 2F). Only 3 DEGs were upregulated in both conditions (Figure 3A), whereas we found 252 genes upregulated in 4 wk HFD but downregulated in 16 wk HFD, mainly linked to cell transport and localisation. Conversely, the 30 genes downregulated in 4 wk HFD, but upregulated in 16 wk HFD referred to immune response (Figure 3A). Finally, a sum of 694 DEGs were downregulated in both 4 and 16 wk HFD, mainly involved in metabolism and transcription (Figure 3A; Table 8). This analysis revealed a large subset of genes being downregulated throughout obesity, but also genes with opposite profile which suggest an adaptive response of CCs to changes in the physiology of the mother. In a parallel approach, we intersected the DEGs datasets from 4wk HFD and 16wk HFD and from the 52 DEGs in common between the two timepoints (Figure 2F) identified 5 main clusters of differently expressed genes. The first 2 clusters comprised 33 genes downregulated in both 4 wk and 16wk HFD (Table 8), with the most significantly deregulated gene at 4 wk HFD *MICAL Like* (*Micall) 1* (FDR = 0.0002), but other genes like *Dynein cytoplasmic 1 heavy chain* (*Dync1h*) or *Collagen* (*Col*) *6a3* (Figure 3B; Table 8) were also found. Clusters 3 and 4 revealed the most interesting set of genes concerning disease progression, due to their opposite profile between 4 wk and 16 wk treatment. In cluster 3 we found genes like *Annexin* (*Anxa*) *11* or *Exportin* (*Xpo*) *5* strongly upregulated in 4 wk HFD, but inhibited in 16 wk HFD (Figure 3B), whereas in cluster 4 we found genes like *Ras homology family member U* (*Rhou*) with opposing profiles at the two time points (Figure 3B). Finally, cluster 5 comprised the 3 genes significantly upregulated throughout obesity (Figure 3B). Therefore, in this analysis we identified gene expression profiles in CCs that represent a valuable tool to assess disease progression in the ovary of obese mice.

**Figure 3.**
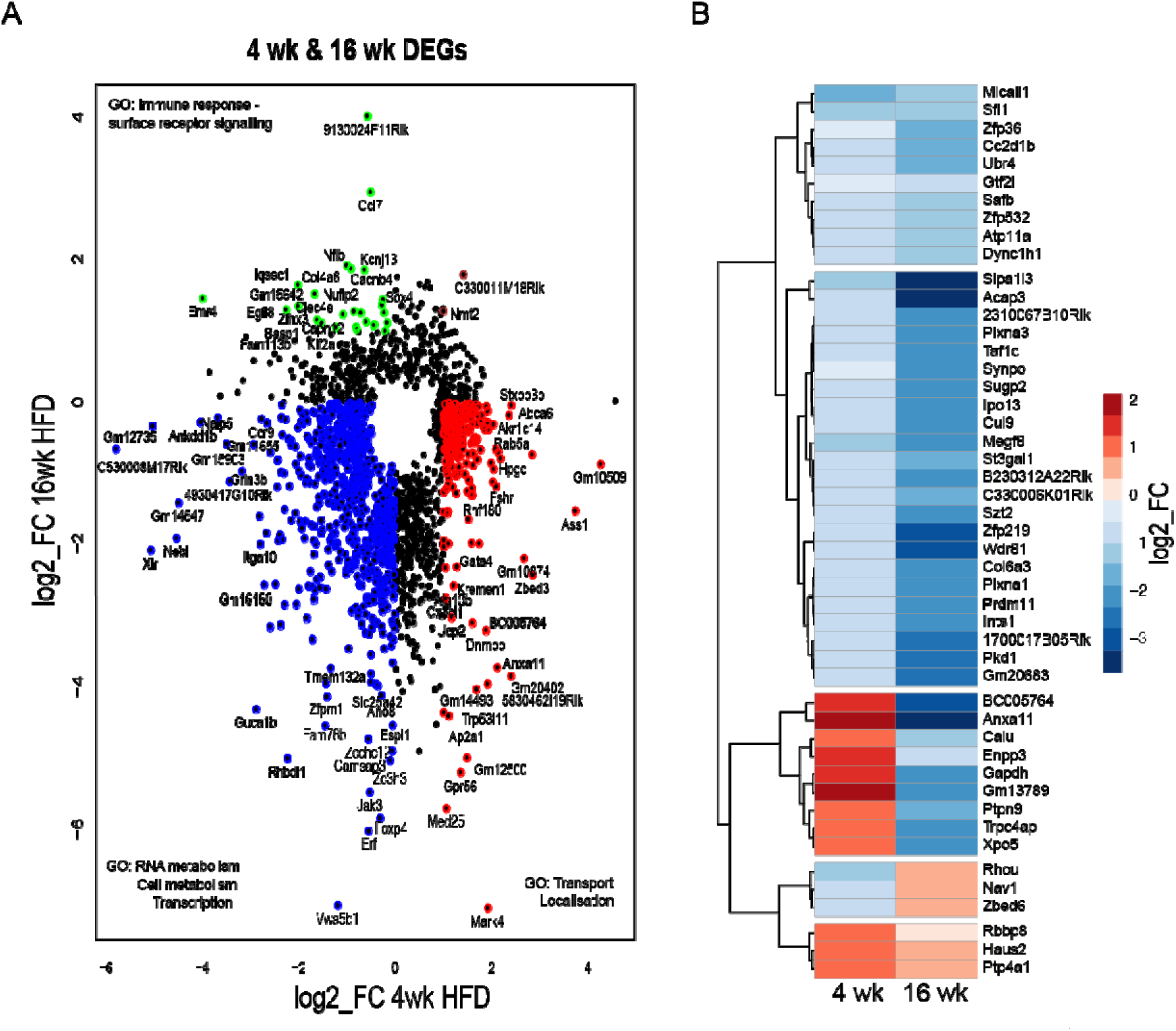
Temporal changes in the transcriptome of cumulus cells during obesity progression. DESeq2 analysis of transcriptome data in cumulus cells (CC) of mice fed high fat diet (HFD) for 4 or 16 weeks (wk). N= 3-7 mice per group. (A) Scatter plot presents genes differentially expressed in at least one condition (False Discovery Rate <0.05) in CC after 4 wk (997 genes) or 16 wk HFD (846 genes). Genes coloured blue are downregulated after both 4wk and 16wk HFD; green downregulated after 4 wk HFD and upregulated after 16wk HFD; brown upregulated after both 4 wk HFD and after 16wk; red upregulated after 4 wk HFD and downregulated after 16 wk HFD. (B) Heatmap presents hierarchical clustering of genes significantly changed in both 4 and 16 wk HFD normalised by average control fed chow diet. Genes cluster in group downregulated after 4 wk and 16 wk HFD, group upregulated after 4 wk and downregulated after 16 wk HFD, group downregulated after 4 wk and upregulated after 16wk HFD and group upregulated after both 4 wk and 16 wk HFD. Gene ontology analysis performed with Gene Ontology Enrichment Analysis and Visualisation Tool. log2_FC of reads per million (RPM).

### 4. The contribution of leptin to changes in gene expression in CCs from obese mice

After identifying the major molecular changes in leptin signalling in the ovaries of DIO females, we aimed to establish an *in vivo* system that would expose the ovaries to the elevated levels of circulating leptin, a feature seen in obesity (7), but lacking all remaining traits of obesity. Thus, we conceived a model for pharmacological hyperleptinemia. Sixteen days of leptin treatment resulted in a consistent drop in BW and FM (Figure supplement 6B, 6E; p<0.01) and increased incidence of oestrus (Figure supplement 6G; p<0.05). The ovaries from animals in oestrus stage were collected for mRNA and protein analysis. Whereas injections of 100 μg leptin for 9 days did not change the protein expression of leptin signalling molecules (Figure supplement 7A-J), ObR expression and phosphorylation of Tyr985 and STAT5 were decreased after 16 d (Figure supplement 7A, 7B, 7G; p<0.05, p=0.09, p<0.01 respectively), together with increased SOCS3 (Figure supplement 7I, 7J; p<0.01, p<0.05 respectively). Validation of a pharmacological hyperleptinemia model allowed us to access ovarian samples from mice with hyperactivation of ObRb, here indicated by increased SOCS3 expression, but lacking the remaining traits of obesity.

Next, we collected CCs from superovulated mice after LEPT treatment and analysed their transcriptome (Figure 4A). After DESeq2 analysis (FDR <0.05), a total of 2026 differently expressed genes were found between LEPT and CONT samples (1212 genes upregulated and 814 downregulated) (Figure 4B). Gene ontology analysis of the DEG lists showed that the upregulated genes for LEPT were associated primarily with cellular organisation, the cytoskeleton and immune responses, supporting the immune-mediating role of leptin as evidenced before (40–42). Conversely, amongst the LEPT downregulated pathways were cell metabolism as well as chromatin organisation and histone modifications (Figure supplement 8; Table 9). Next, based on the hypothesis that early-onset obesity is followed by hyperactivation of leptin signalling in the ovary, we overlapped both 4wk DIO model and LEPT transcriptome datasets, aiming to pinpoint the LEPT driven effects in the CC transcriptome during early obesity. PCA revealed the clustering of 4 wk DIO and CONT samples, and LEPT samples apart (Figure 4C). Indeed, leptin treatment seemed to drive PC1. Next, we overlapped the DEGs from 4 wk HFD and LEPT protocols and found 144 genes upregulated in both LEPT and 4 wk HFD. These were related to response to toxins, transport and glucose metabolism (Figure 4D; Table 10). More specifically, the genes *Lipocalin* (*Lcn*) *2, Anxa11* and *Glucose-6-phosphate dehydrogenase x-linked* (*G6pdx*) were amongst the most significant (Figure 4D). Conversely, the GO terms associated with the 177 downregulated genes in both protocols were metabolism and gene expression regulation (Figure 4D; Table 10). A number of downregulated genes were found to encode important epigenetic factors, such as *Dna segment, chr 14, abbott 1 expressed o* (*Tasor*), *Lysine (k)-specific methyltransferase 2d* (*Kmt2d*/*Mll2*), *Methyl-cpg binding domain protein* (*Mbd*) *2*, and *DNA methyltransferase* (*Dnmt*) *3a* (Figure 4E), which suggested epigenetic dysregulation. Another important effect that could be attributed to leptin in early stages of obesity was the repression of genes mediating actin-cytoskeleton reorganisation (Figure 4F; Table 9). Furthermore, we assessed the potential impact of leptin on genes involved in CC metabolism, and verified the role of leptin on glucose metabolism (Figure 4G) and fatty acid oxidation (Figure 4H), which was reflected in the similarities between LEPT and 4 wk HFD. Here, we questioned how lack of leptin signalling could be detrimental metabolically. For instance, the oocyte is unable to metabolise glucose due to low phosphofructokinase activity (43), highlighting the importance of glycolytic activity of CCs in the generation of pyruvate (44). This function appeared to be decreased in 16 wk HFD, which could be the result of the establishment of leptin resistance in the ovary. As a consequence, the transport of pyruvate into the oocyte would be decreased, which could directly impact the tricarboxylic acid cycle (TCA) and adenosine triphosphate (ATP) generation (Figure supplement 9B) (45). Leptin is also known to be key for free fatty acid (FFA) metabolism, promoting their oxidation and regulating the homeostasis of triglycerides in a cell (46,47). Thus, disruption of leptin signalling in 16 wk HFD CCs (Figure supplement 9A) could be relevant for lipotoxicity and stress previously described in obese ovaries (48) (Figure supplement 9E, 9F). In general, hyperactivation of leptin signalling in CCs seemed to be linked primarily to impaired cell membrane transport and endocytosis, but also cell metabolism and gene expression regulation.

**Figure 4.**
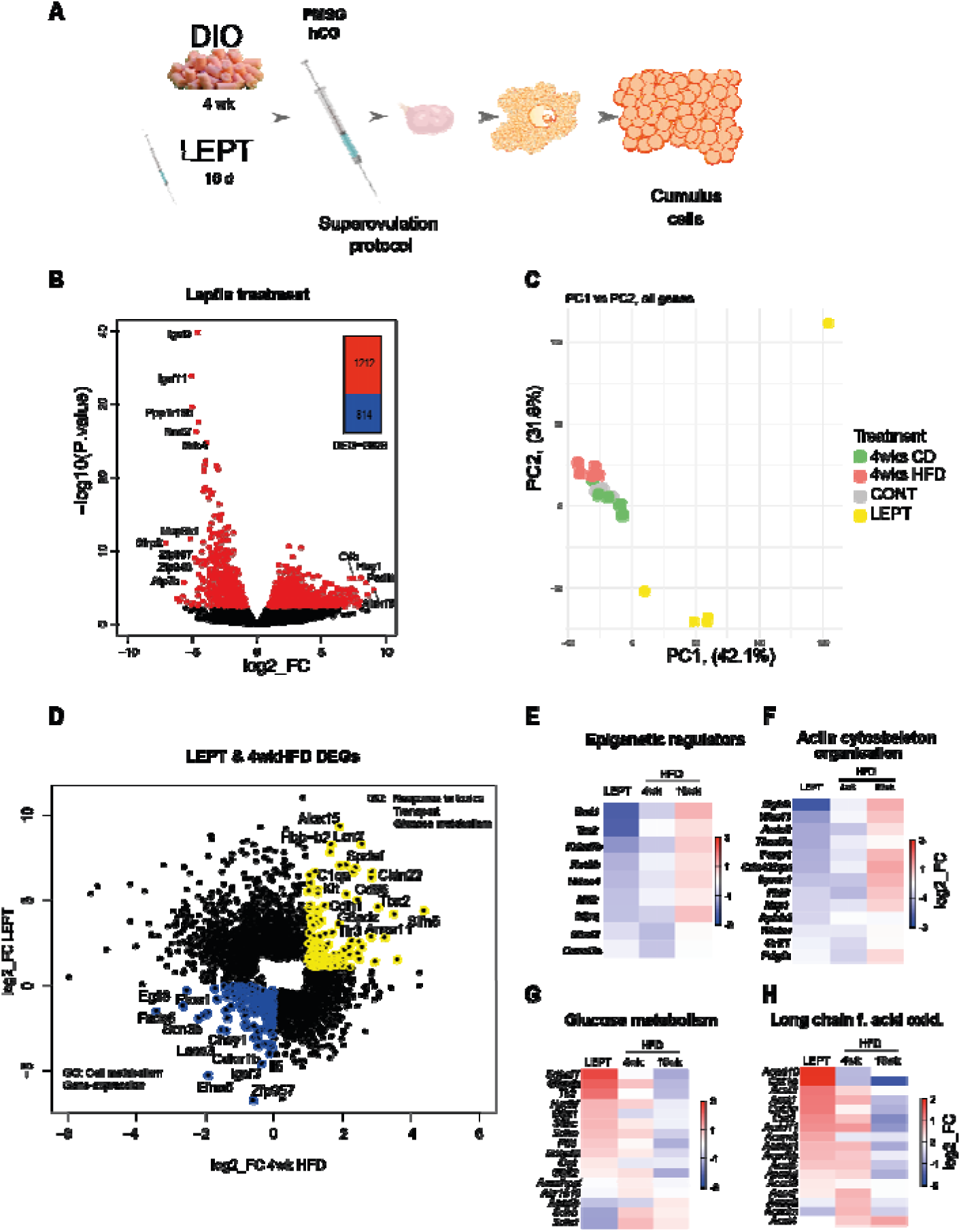
Pharmacologically hyperleptinemic mouse model shows leptin effects in the transcriptome of cumulus cells during early obesity. (A) Experimental design: mice were fed chow diet (CD) of high fat diet (HFD) for 4 weeks (wk) (4 wk DIO) or injected with saline (CONT) or 100μg of leptin (LEPT) for 16 days, followed by superovulation and collection of cumulus cells from cumulus-oophorus-complexes. RNA-seq analysis of gene expression in cumulus cells. N=3-7 mice per group. (B) Volcano plots showing distribution of differentially expressed genes in LEPT group; genes with False Discovery Rate <0.05 coloured red. (C) Principal component analysis of global transcriptome shows LEPT effect is the main source of variance in the data (first principal component, PC1). DESeq2 analysis of transcriptome data in cumulus cells. (D) Scatter plot presents genes differentially expressed in cumulus cells in LEPT or in 4 wk HFD, with False Discovery Rate <0.05. Those coloured blue are down-regulated both in response to leptin treatment and 4 wk HFD; those in yellow upregulated by both treatments. Heatmaps presenting fold of change in expression of genes associated with the following pathways: (E) epigenetic regulation; (F) actin cytoskeleton organisation; (G) glucose metabolism; (H) long chain fatty acid oxidation in CC. Gene ontology analysis performed with Gene Ontology Enrichment Analysis and Visualisation Tool. log2_FC of reads per million (RPM)

## DISCUSSION

The present study characterises the molecular mechanisms underlying the establishment of leptin resistance in the ovary of DIO mice. Furthermore, making use of sensitive methods for reduced-cell number RNA-seq, we studied the transcriptome of the somatic cells surrounding the oocyte from mice subjected to DIO for 4 wk and 16 wk, as well as validated model for pharmacological hyperleptinemia – a system presenting exclusively increased circulating levels of leptin amongst all features of obesity, which allowed us to pinpoint the exclusive effects of leptin-SOCS3 ovarian hyperactivation during early-onset of obesity.

Leptin is a major adipokine, which was initially linked to satiety (6). The establishment of leptin resistance at different levels in the body has been documented in recent years as one of the outcomes of obesity. Accordingly, leptin signalling is deregulated in the hypothalamus (49) and liver of obese human and mice (50). However, the same was not found in kidney (51) or heart (52) of obese humans, suggesting an organ-specific response. We confirmed here, for the first time, the establishment of leptin resistance in the ovary of obese female mice. This poses very important questions on the long-term effects of obesity on ovarian performance, concerning the known local roles of leptin on follicular growth (12), ovulation (13), and oocyte quality (53). Leptin action in the ovary is highly intricate and its effects bimodal. Low leptin levels in circulation facilitate the transition from primary to secondary follicles (12), but leptin is also required for ovulation, possibly supporting CC expansion through cyclooxygenase (COX) 2 and hyaluronic acid synthase (HAS) 2 activity (13). Indeed, *ob/ob* mice contain antral follicles in their ovaries (data not shown), but fail to ovulate. Thus, during obesity progression, altered leptin signalling in the ovary could lead to functional failure and infertility through different mechanisms, mainly characterised by the hyperactivation of ObRb in the onset of obesity and complete failure in signalling in late obesity.

Our first aim was to elucidate the molecular mechanisms leading to the establishment of leptin resistance in the obese ovary. The analysis of different ObRb Tyr domains highlighted the decrease in pTyr985 along with pJAK2 in ovaries from 16 wk HFD mice, concomitant with the increase in SOCS3 protein and decrease in pSTAT5. Functionally, STAT5 phosphorylation in the mouse ovary was shown to be crucial for prolactin signalling and cell proliferation during follicular growth (54), as well as corpus luteum formation (55). Hence, reduced pSTAT5 signalling *per se* could compromise oocyte maturation and fertility during obesity. Importantly, we observed that SOCS3 staining in the oocyte occurred mainly in response to ObRb activation, since the *ob/ob* mouse presented weaker staining. This suggests a direct impact of disrupted ovarian leptin signalling on oocyte quality through SOCS3 activation. Indeed, at 16 wk DIO, we observed different levels of *Socs3* transcribed in various ovarian components. Our RNA-seq data revealed that *Socs3* was increased at 4 wk HFD, but decreased at 16 wk HFD in CCs, whereas in the TC fraction it was upregulated at both time points. This may suggest blunted ObRb signalling in CCs at 16 wks HFD, once the transcription of the major components of the pathway was inhibited (Figure supplement 9A). Therefore, leptin signalling in CCs seems to be highly sensitive to obesity and maternal metabolic performance.

Having an understanding of the impact of obesity on leptin signalling in the ovary, we then analysed the transcriptome of CCs from DIO mice. A major observation of this study was the striking correlation between BW and the global gene expression profile of CCs. On the other hand, other studies showed functional changes in the ovary, including depletion of primordial follicles and inflammation in HFD mice, irrespective of gain in BW (56). Differences in diet composition, as well as variable length of exposure to diet, might account for the differences between studies. Also the aforementioned study did not present a global gene expression analysis. Interestingly, when found that the expression profile of HFD-DEGs in the HFDLG CCs was similar to that in 16 wk CD, clearly demonstrating the impact of maternal BW, which probably largely reflects adiposity in this model, on gene expression in CCs.

Another major outcome of the transcriptome analysis of CCs was the identification of gene signatures altered in early vs late stages of obesity. After 4 wk HFD, mainly genes involved in glucose metabolism and cell membrane trafficking were differently expressed. The use of the pharmacologically hyperleptinemic model allowed us to dissect the contribution of hyperactivation of ObRb to the major changes taking place in CCs in early obesity. Increased activation of the JAK-STAT cascade seemed mainly to impair cellular trafficking and paracrine transfer of macromolecules. This is known to be a crucial process for the metabolic cooperation between the oocyte and somatic cells (23). Cell trafficking and nutrient mobilisation to the oocyte, as well as the uptake of signalling molecules from the oocyte, is fundamental for COCs expansion and oocyte maturation (23). Indeed, the genes *Micall1* and *Unc-51 Like Kinase (Ulk) 4* are important mediators of endocytosis, and were shown to be regulated by *Stat3* (57,58). Furthermore, amongst the genes upregulated in both 4wk HFD and LEPT we found *Lcn2*, associated with lipid and hormone transport (59), *Claudine (Cldn) 22*, a component of tight junctions (60), and *Anxa11*, known to be involved in transmembrane secretion (61). This is suggestive of the effects of leptin in altering transmembrane transport in the early-onset of obesity.

We also identified the metabolic gene *Arachidonate 15-Lipoxygenase* (*Alox15*) and the transcription factor *Hes Related Family BHLH Transcription Factor With YRPW Motif* (*Hey) 1*, were amongst the most significantly upregulated genes in both 4 wk HFD and LEPT (Table 10). The transcriptional repressor HEY1 is directly activated by Notch Receptor (NOTCH) 2 during follicular development, and both HEY1 and NOTCH2 were shown to be increased in proliferating granulosa cells and can contribute to ovarian overstimulation and premature follicular failure (62). These effects further demonstrate the detrimental role of increased ObRb activation during the onset of obesity in cell trafficking and immune response.

Amongst the downregulated signatures in both 4 wk HFD and LEPT we found genes that encoded for important epigenetic factors, such as *Tasor*, *Kmt2d*/*Mll2*, *Mbd2*, and *Dnmt3a* (Figure 4D, table 10), which could indicate epigenetic dysregulation in these cells in early obesity being mediated by leptin. Another striking result was the coordinate downregulation of genes involved in cytoskeleton and actin-filament organisation again in 4 wk HFD and LEPT. As in axons, microtubules form the cytoskeletal core of granulosa cell transzonal projections (TZPs), which provide tracks for the polarized translocation of secretory pathway organelles (63). Thus, by impairing the intrinsic stability of TZPs in granulosa cells, leptin could be affecting the paracrine exchanges between oocytes and somatic cells, an instrumental system for oocyte maturation (64). Indeed, the oocyte is in extreme need of the metabolites generated in CCs, but also signalling factors such as growth differentiation factor (GDF) 9 secreted by the oocyte and required to orchestrate CCs function. Leptin seemed to support the TCA cycle at 4 wk HFD (Figure supplement 9B), which suggested to us that at this early stage the boost in leptin signalling in CCs could actually have beneficial effects, following the positive response on oocyte competence and GDF9 signalling (Figure supplement 9D). However, at 16 wk HFD the inferred drop in CC metabolic fitness was paralleled by a decrease in the main paracrine mediators of oocyte maturation and responsiveness to GDF9 (Figure supplement 9B-D), which invariably suggest compromised oocyte quality. The aforementioned events are an important part of COC expansion, a complex mechanism triggered by luteinizing hormone (LH), in which bidirectional exchange of metabolites and signalling factors between the oocyte and CCs leads to maturation of the gamete and resumption of meiosis (22). This process is tightly regulated by immune mediators, particularly interleukin (IL) 6 (65). Indeed, as well as being highlighted in our transcriptome analysis, the role of leptin in the inflammatory response, in particular mediating innate immunity through IL6, has been described before (55). Consequently, the detrimental effect of obesity could be related to increased leptin signalling at 4 wk HFD, but most likely through its failure at 16 wk HFD (Figure supplement 9A). Generally, in the early stages of obesity, leptin downregulated potentially important epigenetic mediators and genes involved in cytoskeletal organisation in CCs.

The analysis of 16 wk HFD DEGs, as well as the profile of temporal changes revealed genes involved in cell trafficking as *Micall1* or *Dync1h*, involved in protein transport, positioning of cell compartments, and movement of structures within the cell (66) to be decreased in 4 wk and 16 wk HFD. Furthermore, the most increased gene in 16 wk HFD was the *Guanylate-binding protein* (*Gbp*) *8* (Table 7), a component of cellular response to interferon-gamma (67). Another gene upregulated at 16 wk HFD was *Rhou*, a gene that regulates cell morphology (68). Considering also the high expression level of inflammatory mediators at this stage, the activated pathways may well be an outcome of lipotoxicity previously described in the obese ovary (48). Thus, during obesity ovarian cells are trying to accommodate the surplus of lipid compounds, which is likely to activate mechanisms of cellular reorganisation. Overall, early changes in CC transport, gene expression and epigenetic regulation are followed by mounting inflammatory pathways and cellular rearrangement to accommodate the lipid surplus.

In conclusion, we found that the ovaries of obese mice develop leptin resistance and that global gene expression in CCs was strikingly correlated with BW. Mechanistically, failure in ovarian leptin signalling was mediated by SOCS3 overexpression, and inhibition of pTyr985 and pJAK2. Initially, during the onset of obesity the hyperactivation of leptin signalling was linked to increased expression of genes for cell trafficking and cytoskeleton organisation, and inhibition of genes associated with epigenetic regulations in CCs. Conversely, in late obesity, altered gene signatures were mainly linked to inflammatory response and morphological rearrangement (Figure 5). This analysis revealed for the first time the temporal changes in gene expression in CCs during obesity progression. Further studies are being undertaken to understand the impact of these changes in oocyte and early embryo development.

**Figure 5.**
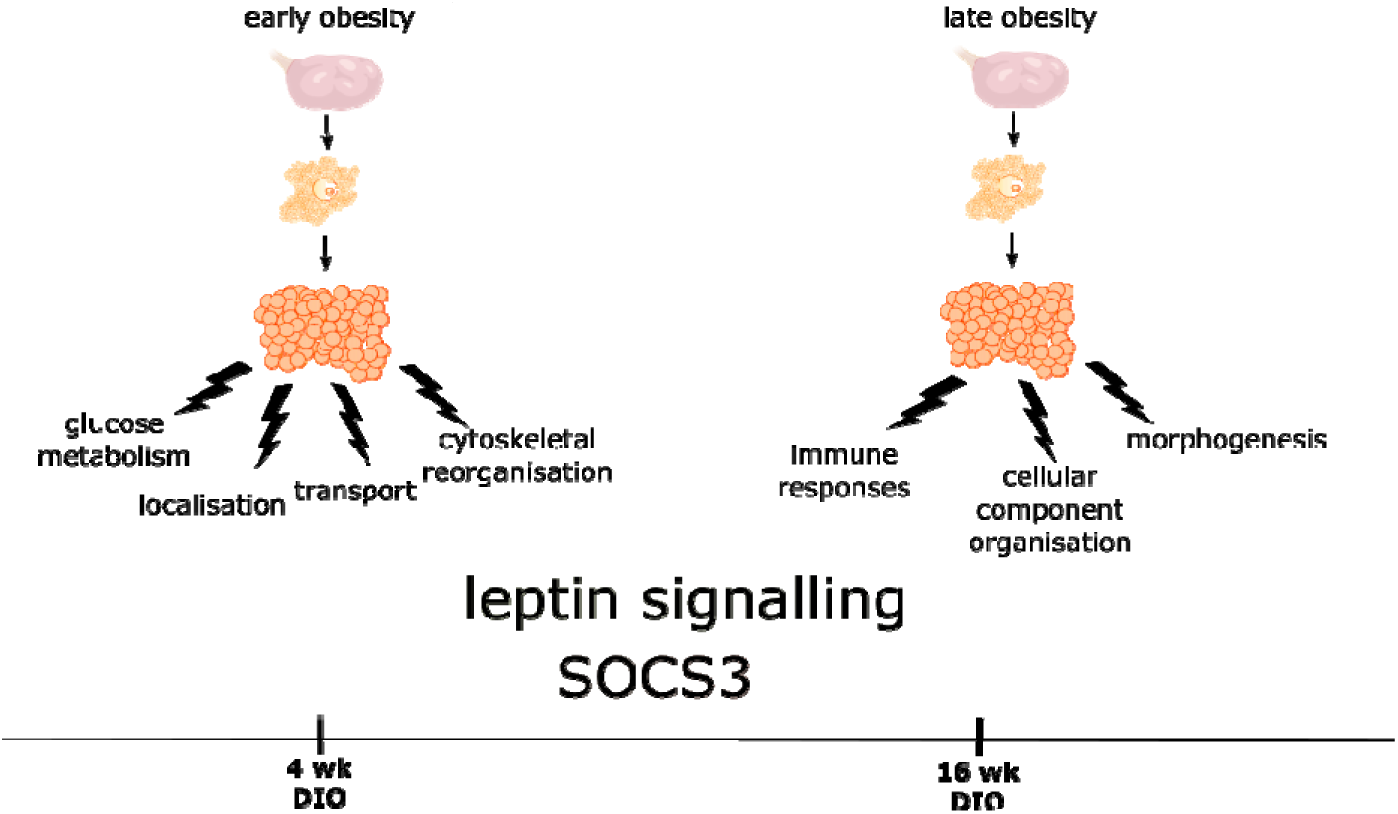
Graphical representation of the main temporal changes in the ovary of obese mice. During early obesity (4 weeks of diet-induced obesity, DIO) increased leptin signalling affects the transcriptome of cumulus cells (CCs). RNA-seq analysis revealed mainly alterations in genes involved in membrane trafficking, cytoskeleton organisation and glucose metabolism. During late obesity (16 wk DIO) leptin resistance is established, which causes accumulation of SOCS3 in the ovary. Transcriptome analysis of CCs at this timepoint indicated the activation of the inflammatory response and cellular anatomical morphogenesis, with inhibition of metabolism and transport.

## AKNOWLEDGEMENTS

We would like to thank Dr Leslie Paul Kozak and Dr Magdalena Jura for their support with the validation and characterisation of the mouse obese phenotype; Dr Jorg Morf for the constructive discussion and suggestions on the analysis of the transcriptome data; Dr Daniel Murta for providing the of mouse ovarian slides; and Dr Fatima Santos and Dr Krzysztof Witek for their support with the imaging and confocal microscopy.

## STATEMENT OF ETHICS

Animal experiments conform to internationally accepted standards and have been approved by the appropriate institutional review body.

## FOUNDING SOURCES

Work was supported by grants from the Polish National Centre for Science (No. 2014/15/D/NZ4/01152 and 2016/23/B/NZ4/03737) awarded to A. G. and the UK Biotechnology and Biological Sciences Research Council and Medical Research Council (BBS/E/B/000C0423, MR/K011332/1, MR/S000437/1) awarded to G.K.; A. G. was supported by Horizon 2020 Marie Curie Individual Fellowship; Processing charge covered by the KNOW Consortium: “Healthy Animal - Safe Food” (Ministry of Sciences and Higher Education; Dec: 05-1/KNOW2/2015).

## AUTHORS CONTRIBUTION

KW data acquisition, analysis and interpretation of the data, writing the manuscript; EW data acquisition, analysis and interpretation of the data; MA, data acquisition and analysis; JCF bioinformatic analysis and interpretation of data, revising the manuscript; GK supervision, funding, revising the manuscript; AG conception and design, funding acquisition, acquisition of data, bioinformatic analysis and interpretation of data, writing and revising of the manuscript.

---

**Figure supplement 1.**
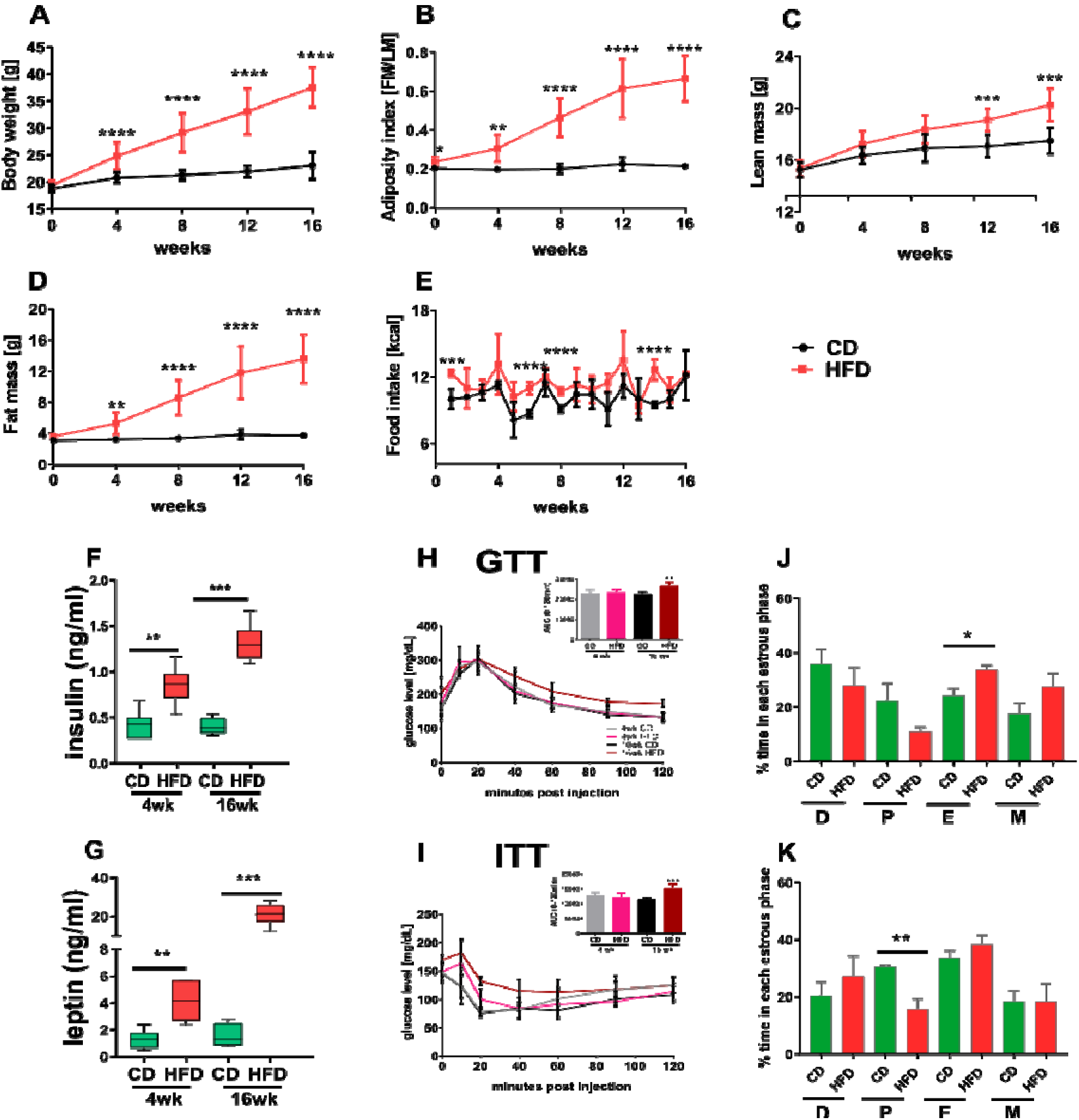
Phenotype characterisation of diet-induced obese mice. Changes in (A) body weight, (B) adiposity index, (C) lean mass, (D) fat mass, (E) food intake in mice fed chow diet (CD, black line) and high fat diet (HFD, red line) for 4 or 16 weeks (wk). Plasma level of (F) insulin and (G) leptin in mice fed CD or HFD for 4 and 16 wk. Glucose tolerance test (GTT, H) and insulin tolerance test (ITT, I) present glucose levels at 0-120 min after glucose and insulin injection, respectively. Bar graphs in the upper right panel present area under the curve for each group. Plasma collected from animals in oestrus phase. Proportion of time spent in each oestrous phase of mice subjected to CD or HFD for 4 wk (J) and 16 wk (K) monitored for 12 days. D, dioestrus; E, oestrus; M, metoestrus; P, pro-oestrus. Each bar represents the mean ± SD for n=12. Differences in phenotype characteristics and plasma hormone level between groups were analysed by Mann-Whitney test, oestrous cycle distribution analysed by unpaired t-test. * p<0.05; ** p<0.01; ***p<0.001.

**Figure supplement 2.**
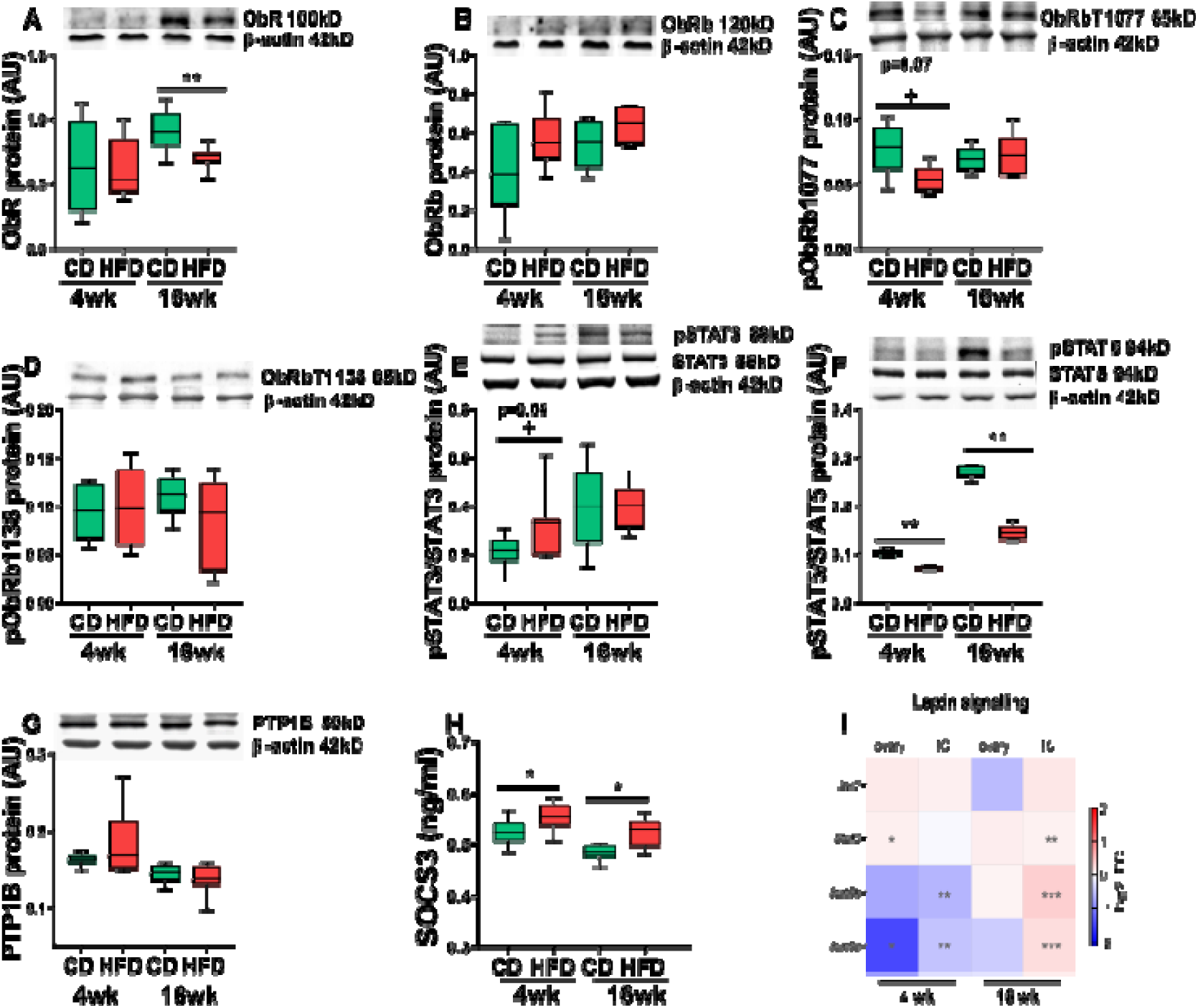
Expression of leptin signalling components in the ovary of diet induced obese mice. Protein abundance of components of the leptin signalling pathway in ovarian extracts analysed by Western blot or ELISA. Animals were maintained on chow diet (CD) or high fat diet (HFD) for 4 or 16 weeks (wk). Abundance of (A) leptin receptor (ObR), phosphorylation of (B) long isoform of leptin receptor (ObRb), (C) tyrosine 1077 of leptin receptor, (D) tyrosine 1138 of leptin receptor, (E) STAT3, (F) STAT5, expression of (G) PTP1B. (H) SOCS3 ovarian quantification in ELISA test. (I) Heatmap showing fold of change in expression of mRNA of leptin signalling components measured in whole ovary or theca/stroma enriched (TC) fraction by RT-PCR. mRNA expression of *Rpl37* and protein expression of β-actin were used to normalize the expression data. Each bar represents the mean ± SD. Differences between groups were analysed by Mann-Whitney test. N=4-8 for immunoblots and N=8 for RT-PCR analysis and ELISA. * p<0.05; ** p<0.01; ***p<0.001; + p=0.06 or p=0.07.

**Figure supplement 3.**
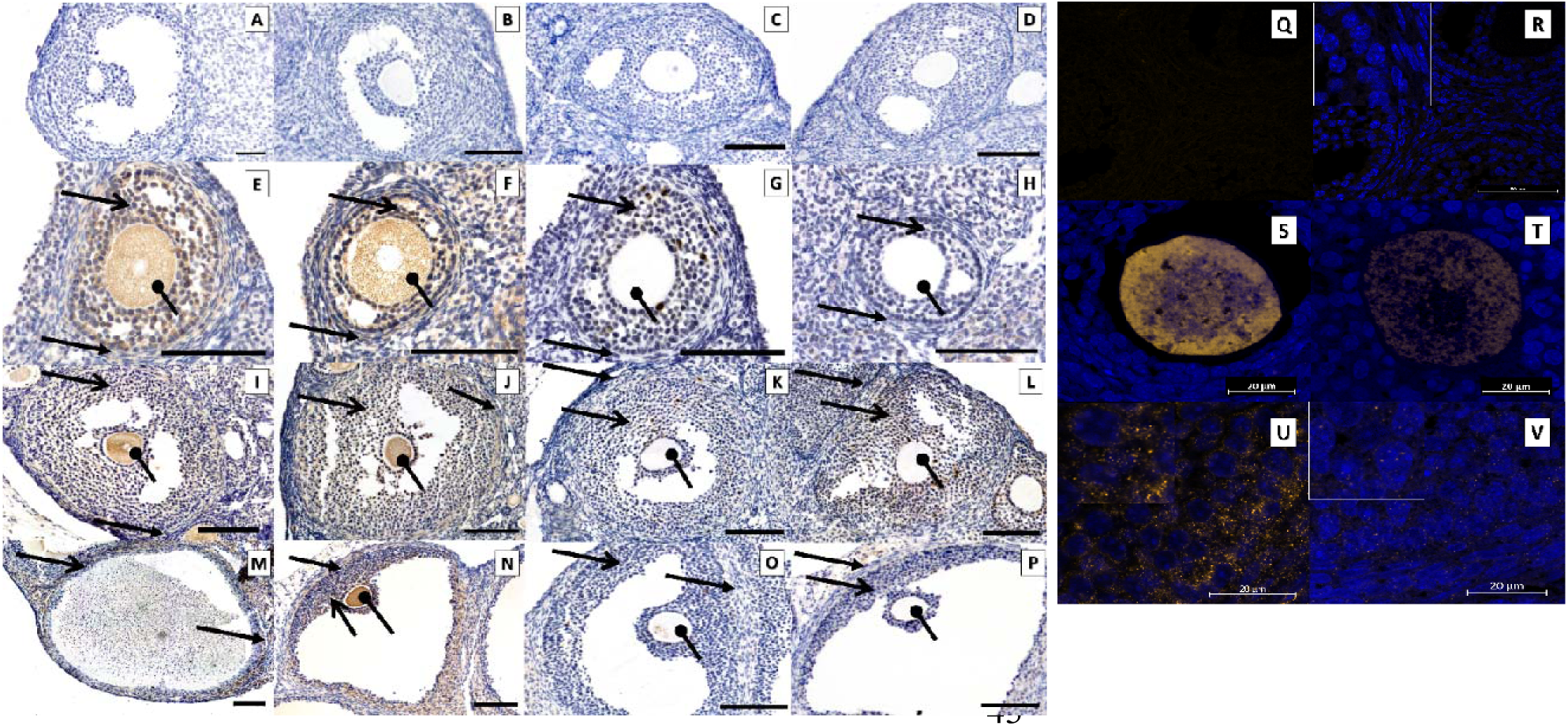
Immunolocalisation of SOCS3 and PTP1B protein in the ovary. Immunohistochemical localisation of SOCS3 and PTP1B protein during follicle development in mice fed chow diet (CD) or high fat diet (HFD) for 4 and 16 weeks (wk) and intraperitoneally injected with saline (C) or leptin (L) for 16 days (d). Positive staining in brown, counterstaining with heamatoxylin. Negative control stained with polyclonal rabbit IgG (A, B) 4 wk CD, (C, D) 4 wk HFD, localisation in secondary follicle SOCS3 (E) 4 wk CD and (F) 4 wk HFD, PTP1B (G) 4 wks CD, (H) 4 wks HFD, antral follicles SOCS3 (I) 16 wk CD, (J) 16 wk HFD, PTP1B (K) 16 wk CD, (L) 16 wk HFD, preovulatory follicle SOCS3 (M) 16 C, (N) 16 L, PTP1B (O) 16 C, (P) 16 L. The scale bar represents 100μm. The specificity of SOCS3 staining was confirmed by immunofluorescent localisation in *ob/ob* mice with genetic deficiency of leptin. Positive staining in orange, nuclear counterstaining with DAPI in blue. (Q-R) negative control 16 wk CD performed with polyclonal rabbit IgG, SOCS3 localised in (S, T) secondary follicle and (U,V) antral follicle from controls (*ob/ob. +/+*; S,U) and leptin deficient ovaries (*ob/ob -/-*; T,V). Images are representatives of 3 biological replicates. Inserts in left top corners are the amplifications of granulosa cells. Pictures are representatives of 3 biological replicates. The scale bar represents 20μm.

**Figure supplement 4.**
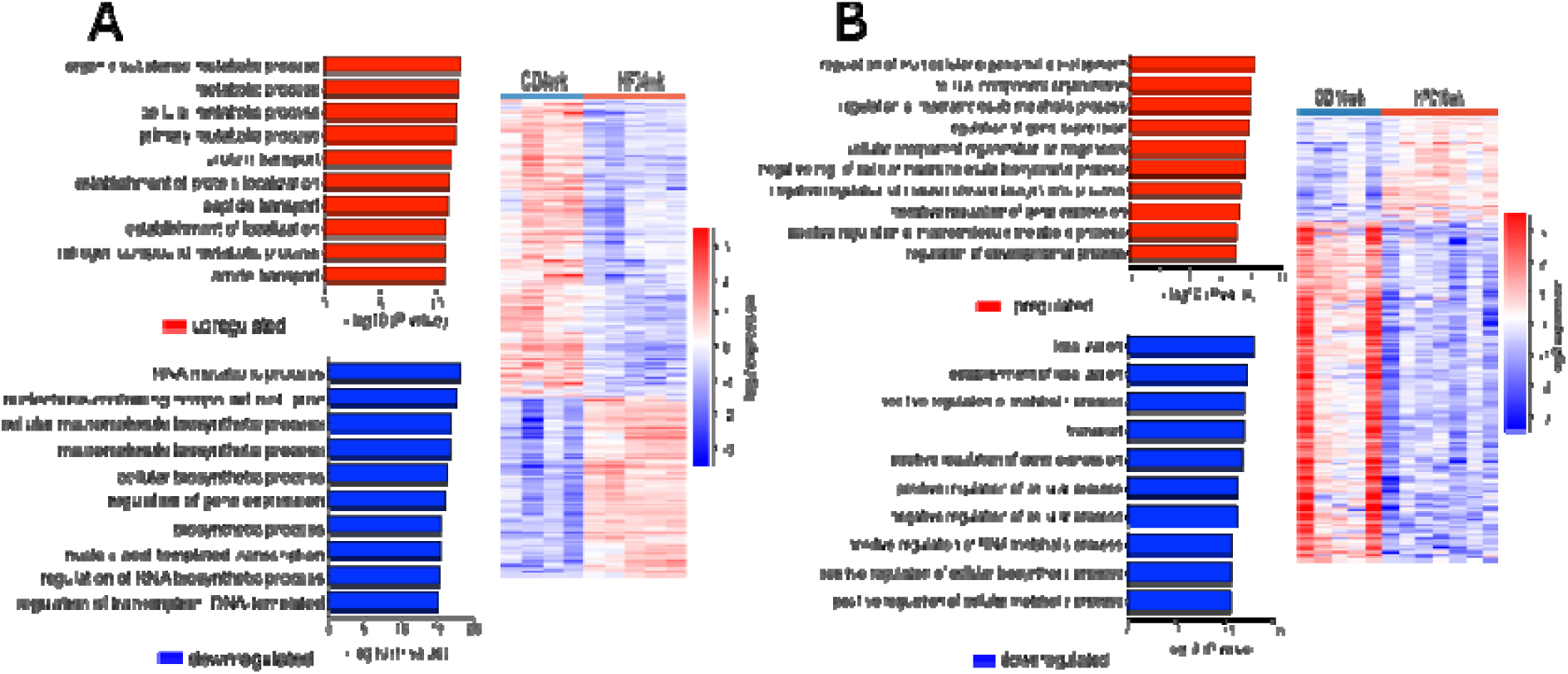
Differentially expressed genes and associated pathways in cumulus cells from diet-induced obesity protocol. DESeq2 analysis of transcriptome data in cumulus cells obtained from mice after 4 or 16 weeks (wk) of chow diet (CD) or high fat diet (HFD). N= 3-7 mice per group. On the right heatmap showing hierarchical clustering of (A) 997 differentially expressed genes after submitting mice to 4 weeks of CD and HFD, (B) 846 differentially expressed genes after submitting mice to 16 weeks of CD and HFD. On the left presentation of pathways of genes with the most significant enrichment after gene ontology analysis. Gene ontology analysis performed with Gene Ontology Enrichment Analysis and Visualisation Tool.

**Figure supplement 5.**
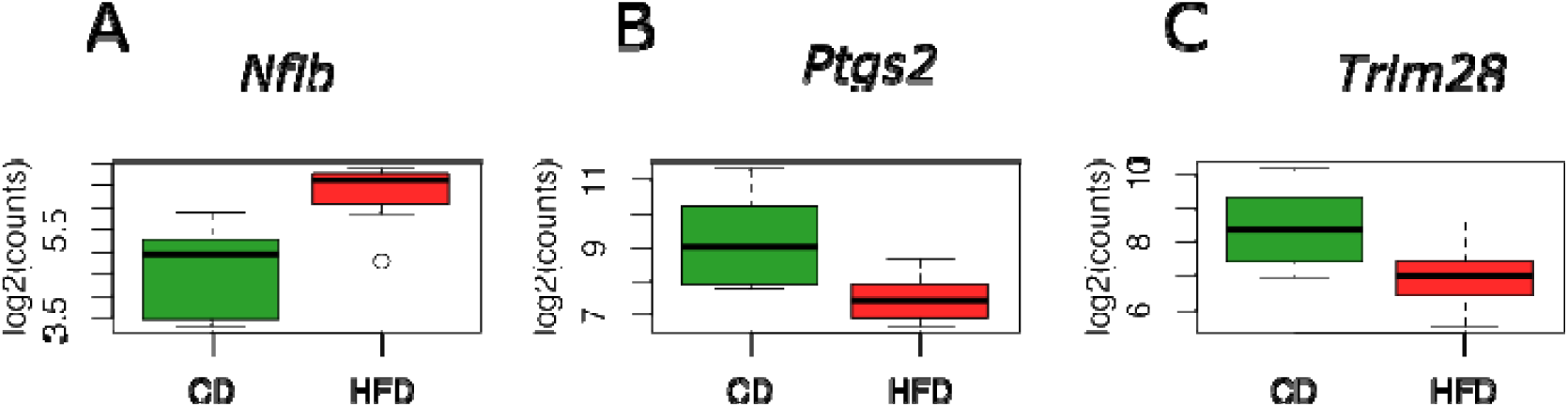
Oocyte competence and embryo quality markers differentially expressed in cumulus cells from mice with late obesity. DESeq2 analysis of transcriptome data in cumulus cells obtained from mice after 16 weeks of chow diet (CD) or high fat diet (HFD). N= 3-7 mice per group. Expression of embryo quality markers (A) *nuclear factor I B (Nfib)*, (B) *cyclooxygenase 2 (Ptgs2)* and oocyte competence marker (C) *tripartite motif containing 28 (Trim28)*. Log2 of counts.

**Figure supplement 6.**
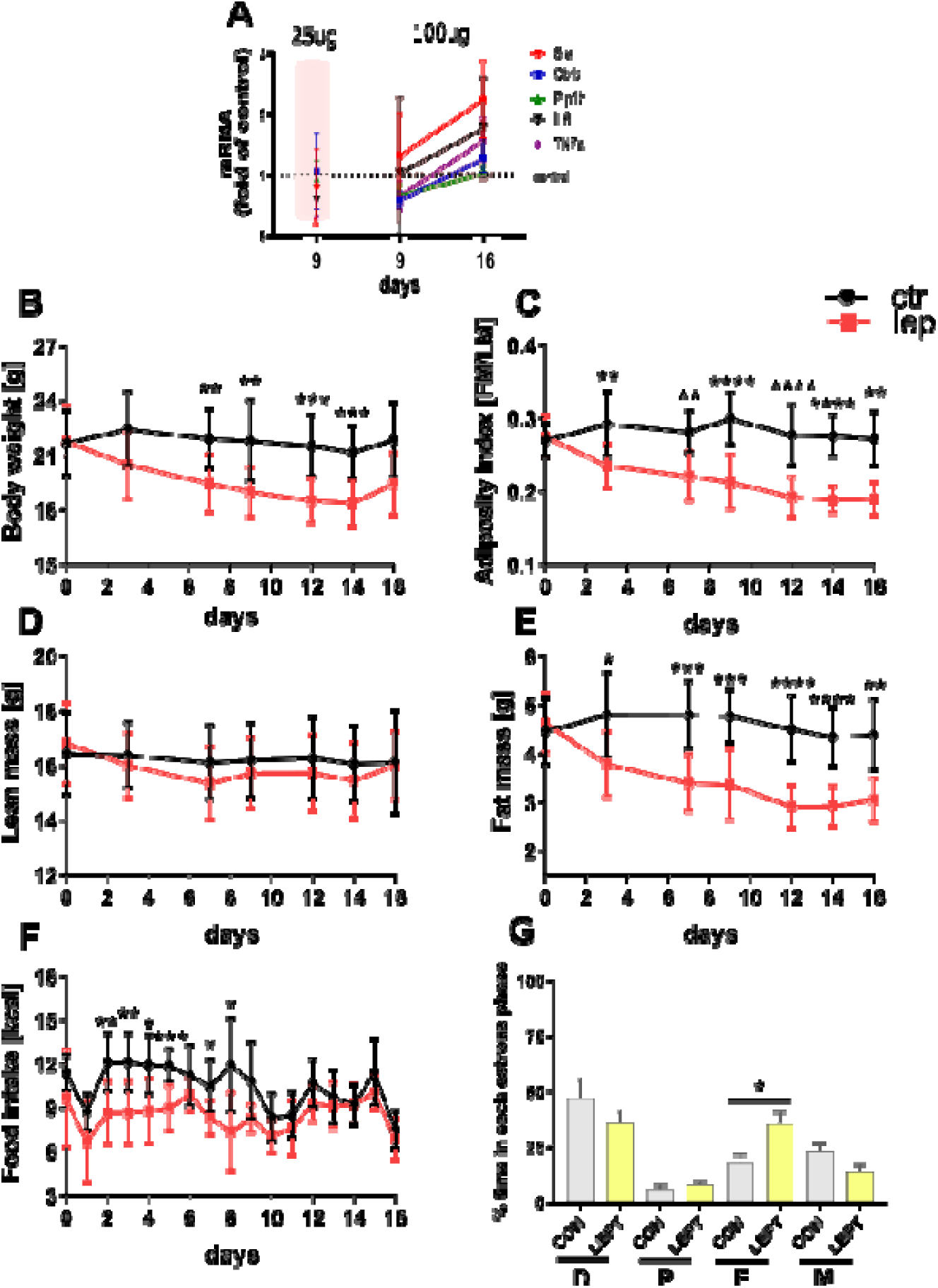
Pharmacologically hyperleptinemic mouse model validation. In order to validate the length of the treatment and the dose of leptin, we analysed whole ovary mRNA from animals treated with 25 or 100 μg of leptin for 9 or 16 d. Injection of 100 μg for 16 days caused changes in the abundance of leptin-responsive transcripts (69–72) in ovarian extracts collected from animals in oestrous stage. Animals were injected with saline (C) or different doses of leptin (L) for 9 or 16 days (d). (A) mRNA level of *steroidogenic acute regulatory protein (Star), long isoform of leptin receptor Obrb, protein tyrosine phosphatase non-receptor type 1 (Ptp1b), interleukin 6 (Il6), tumor necrosis factor a (Tnfa)* expressed as fold of control after injecting animals for 9 or 16 days with 25μg or 100 μg of leptin. Changes in (B) body weight, (C) adiposity index, (D) lean mass, (E) fat mass, (F) food intake in mice intraperitoneally injected with saline (ctr, black line) or leptin (lep, red line) for 16 days. (G) Proportion of time spend in each oestrous phase of hyperleptinemic mice. D, dioestrus; E, oestrus; M, metoestrus; P, pro-oestrus. Each bar represents the mean ± SD. Differences in phenotype characteristics between groups were analysed by Mann-Whitney test, oestrous cycle distribution analysed by unpaired t-test. Data show mean values for n=10. * p<0.05; ** p<0.01; ***p<0.001

**Figure supplement 7.**
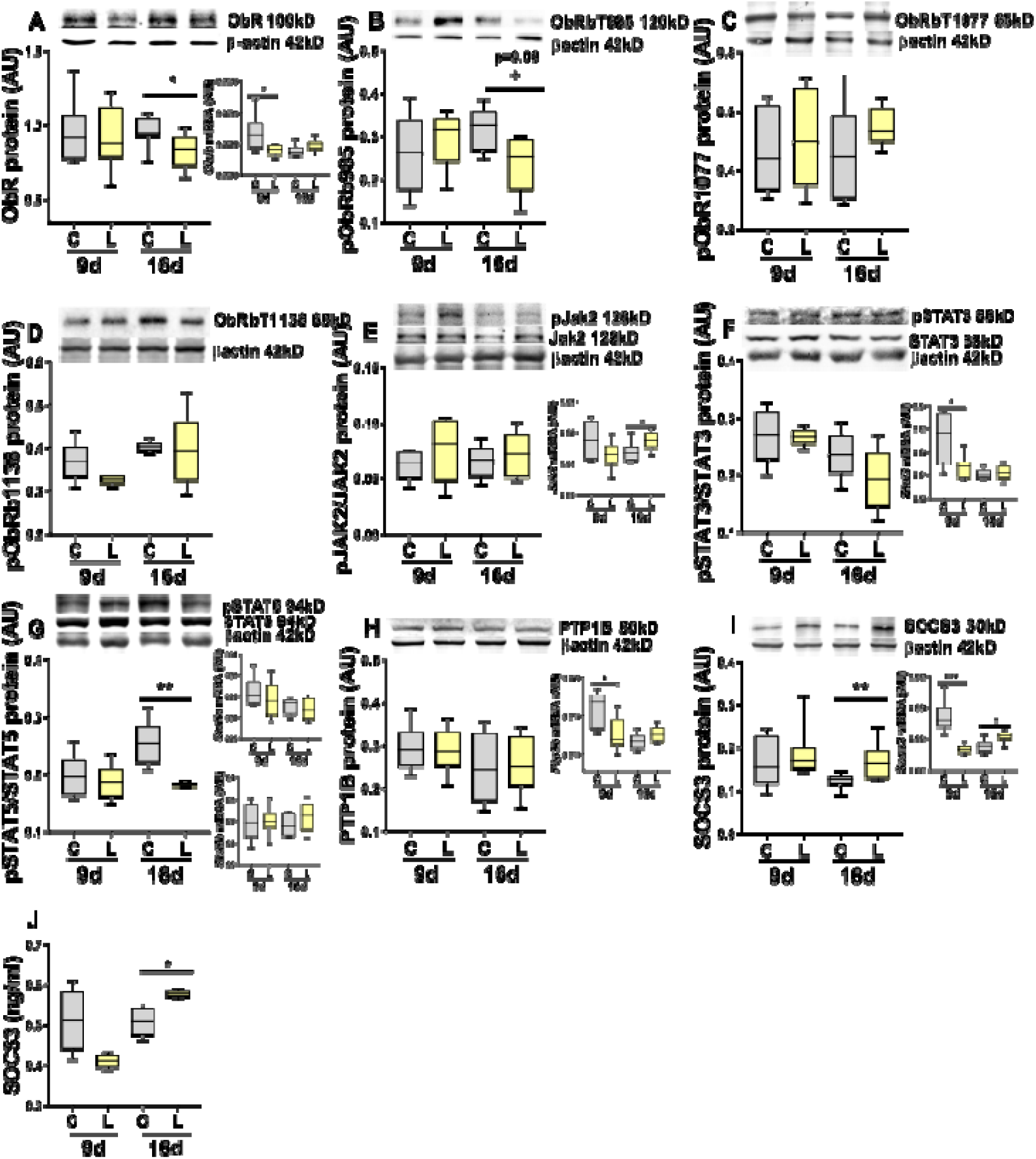
Expression of leptin signalling components in the ovary of pharmacologically hyperleptinemic mice. Abundance of mRNA (grey box) and protein of leptin signalling pathway components in ovarian extracts collected from animals injected with saline (C) or 100 μg of leptin (L) for 9 or 16 days (d) and sacrificed in oestrus stage. Expression of (A) leptin receptor (ObR), phosphorylation of (B) tyrosine 985 of leptin receptor, (C) tyrosine 1077 of leptin receptor, (D) tyrosine 1138 of leptin receptor, (E) Janus kinase 2 (JAK2), (F) signal transducer and activator of transcription 3 (STAT3), (G) STAT5, expression of (H) protein tyrosine phosphatase 1B (PTP1B) and (I) suppressor of cytokine signaling 3 (SOCS3) determined by real-time PCR and Western blot. (J) SOCS3 ovarian quantification in animals in oestrus stage determined by ELISA. mRNA expression of *Rpl37* and protein expression of β-actin was used to normalize the expression data. Each bar represents the mean ± SD. Differences between groups were analysed by Mann-Whitney test. N=4-8 for immunoblots and N=8 for RT PCR analysis and ELISA. * p<0.05; ** p<0.01; ***p<0.001; + p=0.09.

**Figure supplement 8.**
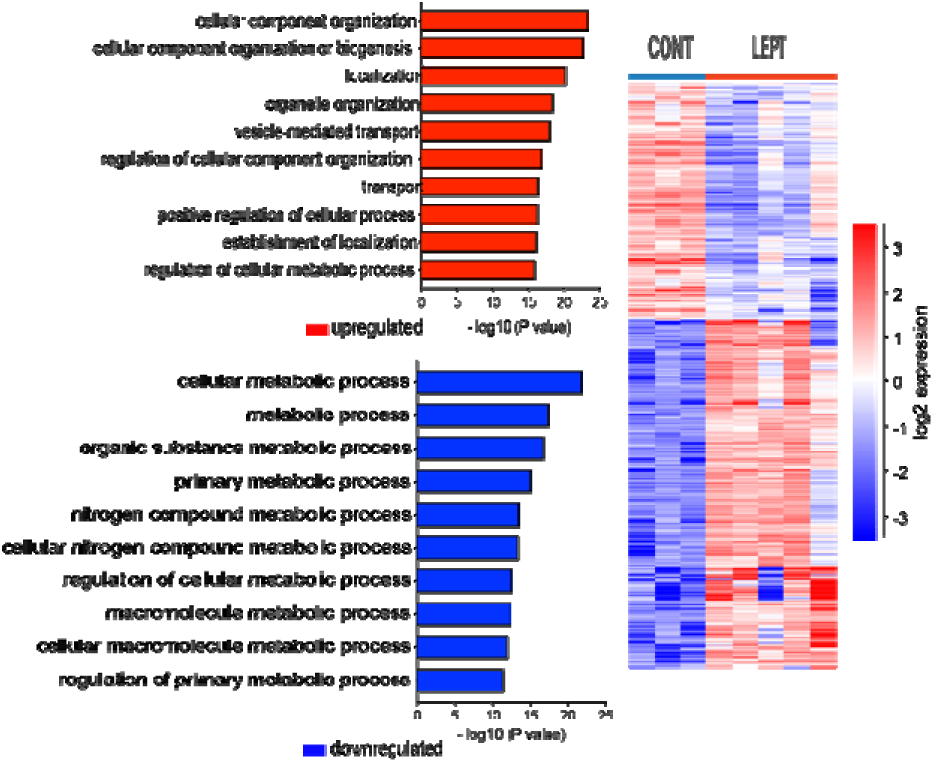
Differentially expressed genes and associated pathways in cumulus cells from hyperleptinemic mice. DESeq2 analysis of transcriptome data in cumulus cells obtained from mice treated with saline (CONT) and leptin (LEPT). N= 3-7 mice per group. On the right heatmap showing hierarchical clustering of 2026 differentially expressed genes after treating mice with leptin for 16 days. On the left presentation of pathways of genes with the most significant relevance after gene ontology analysis. Gene ontology analysis performed with Gene Ontology Enrichment Analysis and Visualisation Tool.

**Figure supplement 9.**
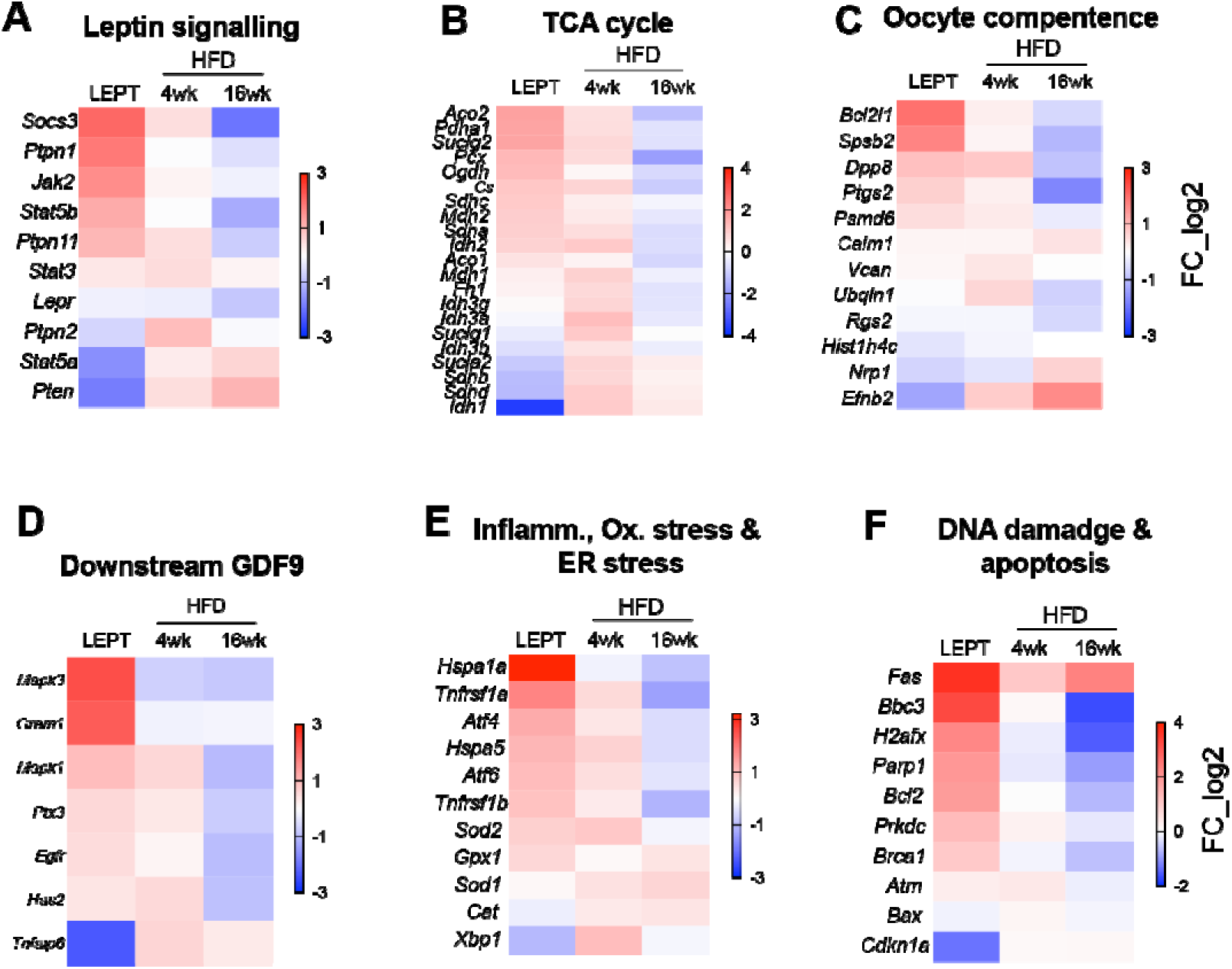
Similarities between profiles of genes differentially expressed in cumulus cells in diet induced-obese mice and leptin treated mice. DESeq2 analysis of transcriptome data in cumulus cells (CC) obtained from mice treated with saline (CONT) and leptin (LEPT) or after 4 or 16 weeks (wk) of chow diet (CD) or high fat diet (HFD). N= 3-7 mice per group. Heatmaps representing fold change in expression of genes associated with the following pathways or processes: (A) leptin signalling, (B) tricarboxylic acid (TCA) cycle, (C) oocyte competence, (D) genes regulated by oocyte-derived growth differentiation factor (GDF) 9, (E) inflammation, oxidative stress and endoplasmic reticulum stress, (F) DNA damage and apoptosis in CC. log2_FC of reads per million (RPM)

