## Supplemental Table 1 for "Leptin Resistance in the Ovary of Obese Mice Is Associated with Profound Changes in the Transcriptome of Cumulus Cells"

Supplementary file 1 Table 1. Specific primer sequences used for quantitative real-time PCR.

| Functional pathway | Gene name | Gene symbol | GeneBank Accession no. | Sequences 5’-3’ | Length (base pair) |
|---|---|---|---|---|---|
| Leptin signalling | Leptin receptor | *Obrb* | NM_146146.2 | CCTCCAGGAGAGATGCTCACAC<br>TGACTGTGCGTGGAACAGGT | 111 |
| | Janus kinase 2 | *Jak2* | NM_001048177.**2** | GGTGTTTCACAAAATCAGGAATG<br>TGTGCAGTTGACCATAATCTCC | 119 |
| | SH2B adaptor protein 1 | *Sh2b* | NM_001289539.**1** | GGACCCAGCGAGAGTAACGA<br>GCAGCAATGGAGGCAGAACT | 101 |
| | Signal transducer and activator of transcription 3 | *Stat3* | NM_011486.5 | CGATGCCTGTGGGAAGAGTC<br>CTGTCACTACGGCGGCTGTT | 97 |
| | Signal transducer and activator of transcription 5A | *Stat5a* | NM_001164062.1 | GGGACAATGCCTTTGCTGAG<br>AGCCCCGGTTGCTCTGTACT | 123 |
| | Signal transducer and activator of transcription 5B | *Stat5b* | NM_001113563.2 | CAGGACAACAATGCCACAGC<br>TTTGGCCGATCAGGAAACAC | 172 |
| | Protein tyrosine phosphatase, non-receptor type 2 | *Ptpn2* | NM_008977.3 | CGCTCTGGCACCTTCTCTCT<br>GGAAAGGCAGGATCTCTCGA | 283 |
| | Protein tyrosine phosphatase non receptor type 1 | *Ptp1b* | NM_011201.3 | TGGCCACAGCAAGAAGAAAA<br>GGAAAGGCAGGATCTCTCGA | 151 |
| | Suppressor of cytokine signaling 3 | *Socs3* | NM_007707.3 | GCGAGAAGATTCCGCTGGTA<br>TACTGATCCAGGAACTCCCGA | 151 |
| Reference genes | Ribosomal protein L37 | *Rpl37* | NM_026069.3 | CTGGTCGGATGAGGCACCTA<br>AAGAACTGGATGCTGCGACA | 108 |
| | Eukaryotic translation initiation factor 5A | *Eif5a* | NM_001166594.1 | CCTCAGCCACCTTCCCAAT<br>AAATGTCAATGCCAACCAGATG | 150 |
