## Supplemental Table 2 for "Leptin Resistance in the Ovary of Obese Mice Is Associated with Profound Changes in the Transcriptome of Cumulus Cells"

Supplementary file 2 Table 2. Specification of antibodies used for Western blot.

| Antibody name and specificity | Company, Cat no, RRID no | Antibody dilution |
|---|---|---|
| Mouse monoclonal against leptin receptor (ObR) | Santa Cruz Biotechnology Cat# sc-8391, RRID:AB_627882 | 1:500 |
| Goat polyclonal against phosphorylated Tyr-985 of leptin receptor (p-ObR Tyr985) | Santa Cruz Biotechnology Cat# sc-16419, RRID:AB_2234640 | 1:500 |
| Rabbit polyclonal against phosphorylated Tyr 1077 ObR (p-ObR Tyr1077) | Millipore Cat# 07-1317, RRID:AB_1977322 | 1:500 |
| Goat polyclonal against phosphorylated Tyr 1138 ObR (p-ObR Tyr1138) | Santa Cruz Biotechnology Cat# sc-16421, RRID:AB_2288076 | 1:500 |
| Rabbit polyclonal against Janus kinase 2 (Jak2) | Santa Cruz Biotechnology Cat# sc-294, RRID:AB_631854 | 1:200 |
| Rabbit polyclonal against phosphorylated Tyr 1007/1008 Jak2 (pJak2) | Santa Cruz Biotechnology Cat# sc-16566-R, RRID:AB_653287 | 1:200 |
| Rabbit polyclonal against Signal transducer and activator of transcription 3 (STAT3) | Santa Cruz Biotechnology Cat# sc-482, RRID:AB_632440 | 1:200 |
| Mouse monoclonal against phosphorylated Tyr 705 Signal transducer and activator of transcription 3 (pSTAT3) | Santa Cruz Biotechnology Cat# sc-8059, RRID:AB_628292 | 1:200 |
| Rabbit polyclonal against Signal transducer and activator of transcription 5 (STAT5) | Santa Cruz Biotechnology Cat# sc-835, RRID:AB_632446 | 1:200 |
| Mouse monoclonal against phosphorylated Tyr 694/699 Signal transducer and activator of transcription 5 (pSTAT5) | Santa Cruz Biotechnology Cat# sc-81524, RRID:AB_1129712 | 1:200 |
| Goat polyclonal against Protein tyrosine phosphatase 1B (PTP1B) | Santa Cruz Biotechnology Cat# sc-1718, RRID:AB_2174942 | 1:200 |
| Mouse monoclonal against Suppressor of cytokine signalling 3 (SOCS3) | Santa Cruz Biotechnology Cat# sc-51699, RRID:AB_630243 | 1:500 |

| | | |
|---|---|---|
| Mouse monoclonal against β-actin | Sigma Aldrich Cat# A2228, RRID:AB_476697 | 1:10000 |
